# Bacteriophage λ RexA and RexB Functions Assist the Transition from Lysogeny to Lytic Growth

**DOI:** 10.1101/2021.04.19.439022

**Authors:** Lynn C. Thomason, Carl J. Schiltz, Carolyn Court, Christopher J. Hosford, Myfanwy C. Adams, Joshua S. Chappie, Donald L. Court

## Abstract

The CI and Cro repressors of bacteriophage λ create a bistable switch between lysogenic and lytic growth. In λ lysogens, CI repressor expressed from the *P*_RM_ promoter blocks expression of the lytic promoters *P*_L_ and *P*_R_ to allow stable maintenance of the lysogenic state. When lysogens are induced, CI repressor is inactivated and Cro repressor is expressed from the lytic *P*_R_ promoter. Cro repressor blocks *P*_RM_ transcription and CI repressor synthesis to ensure that the lytic state proceeds. RexA and RexB proteins, like CI, are expressed from the *P*_RM_ promoter in λ lysogens; RexB is also expressed from a second promoter, *P*_LIT_, embedded in *rexA.* Here we show that RexA binds CI repressor and assists the transition from lysogenic to lytic growth, using both intact lysogens and defective prophages with reporter genes under control of the lytic *P*_L_ and *P*_R_ promoters. Once lytic growth begins, if the bistable switch does return to the immune state, RexA expression lessens the probability that it will remain there, thus stabilizing the lytic state and activation of the lytic *P*_L_ and *P*_R_ promoters. RexB modulates the effect of RexA and may also help establish phage DNA replication as lytic growth ensues.

## Introduction

λ is a temperate bacteriophage and thus can exist in either the lysogenic or lytic state (H Echols, 1986; Oppenheim, Kobiler, Stavans, Court, & Adhya, 2005). The transition between these two states is mediated by two λ regulators: CI and Cro (Eisen et al., 1975; Ptashne & Hopkins, 1968). In the lysogenic state, the phage chromosome is integrated into that of its host *Escherichia coli* and is quiescent with its lytic functions repressed by the CI protein (Ptashne & Hopkins, 1968). In the repressed lysogen, pairs of CI dimers bind cooperatively to operator sites *O*_L1_ and *O* _L2_ on the left and *O*_R1_ and *O* _R2_ on the right to repress the early *P*_L_ and *P*_R_ promoters (Figure 1), respectively, and thereby, inhibit transcription of all the lytic genes downstream from those promoters (A. D. Johnson, Meyer, & Ptashne, 1979). Interaction between CI repressor molecules bound to the left and right operators results in topological looping of the intervening DNA, which contains the phage immunity region (Dodd, Shearwin, & Egan, 2005); this looping enhances repression of *P*_L_ and *P*_R_ and stabilizes the lysogenic state. In a lysogen, the *c*I repressor gene is transcribed and expressed from the *P*_RM_ promoter (Spiegelman et al., 1972). When CI repressor is bound to *O* _R1_ *O* _R2_, it activates transcription initiation by RNA polymerase at the *P*_RM_ promotor (Lewis, Gussin, & Adhya, 2016). Two other genes, *rexA* and *rexB*, are downstream of the *c*I gene and, like *cI*, are expressed as part of the *P*_RM_ operon (Benzer, 1955; Matz, Schmandt, & Gussin, 1982) (Figure 1). A second promoter, *P*_LIT_, is located within the distal end of *rexA*, and transcribes just *rexB* (Hayes, Bull, & Tulloch, 1997; Landsmann, Kroger, & Hobom, 1982). Thus, *rexB* can be transcribed by either *P*_RM_ and/or *P*_LIT_ (Hayes & Szybalski, 1973; Liu, Jiang, Gu, & Roberts, 2013; Thomason et al., 2019). A λ lysogen can switch to the lytic pathway, produce progeny phage, and lyse its host, releasing phage. This can happen in response to DNA damage, which in *E. coli* initiates the SOS response (d’Ari, 1985; Witkin, 1991). Single-stranded DNA is generated as a result of the damage and is bound by the RecA protein, converting RecA to an activated DNA-bound form, RecA*. RecA* binds the CI repressor and acts as a co-protease, initiating auto-cleavage of the CI protein (Craig & Roberts, 1980; Ennis, Ossanna, & Mount, 1989). Loss of CI repression leads to activation of the lytic *P_L_* and *P_R_* promoters. The *cro* gene, which encodes the Cro repressor, is the first transcribed gene from the *P_R_* promoter (H. Echols, Green, Oppenheim, Oppenheim, & Honigman, 1973; Folkmanis, Maltzman, Mellon, Skalka, & Echols, 1977). Cro repressor binds to the same operator sites as CI, but with a different pattern of affinities (Darling, Holt, & Ackers, 2000; A. Johnson, Meyer, & Ptashne, 1978). As Cro repressor is made, it first binds *O*_R3_ and represses the pRM promoter, thus blocking *P*_RM_ and *c*I gene transcription, and further reducing CI repressor levels and stabilizing the switch to lytic expression. Thus, Cro locks in the anti-CI repressor state, also called the nonimmune state, ensuring lytic growth (Ptashne et al., 1980; Svenningsen, Costantino, Court, & Adhya, 2005).

**Figure 1.**
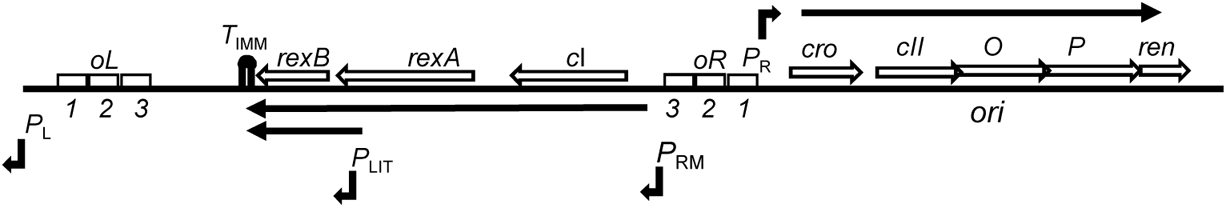
Genetic map of λ immunity region and DNA replication genes. The CI and Cro repressors are expressed in the *P*_RM_ and *P*_R_ operons, respectively, and are major players in the lysis-lysogeny decision. These two repressors control expression of the major lytic promoters *P*_L_ and *P*_R_ by binding to the left and right tripartite operator sites, *O*_L_ and *O*_R_, with the lytic *P*_R_ and lysogenic *P*_RM_ promoters sharing coordinate but opposing regulation within the *O*_R_ segment. The *rexA* and *rexB* genes are downstream of *c*I. The *P*_LIT_ promoter is embedded in the terminal coding sequence of *rexA*, with the consequence that *rexB* is transcribed from two promoters, *P*_RM_ and *P*_LIT_, while *c*I and *rexA* are transcribed only from *P*_RM_ (Thomason et al 2019). Black arrows represent the beginning of the various promoter transcripts shown. Transcription from *P*_RM_ and *P*_LIT_ ends at the transcriptional terminator, *T*_IMM_, immediately downstream of *rexB*. The DNA replication genes *O* and *P* are transcribed from *P*_R_, as is *ren*, which is also likely involved in DNA replication.

λ *c*I^+^ lysogens, which express wild type CI repressor, are maintained in a repressed state at any growth temperature. A mutation in the *c*I gene, *c*I*857*, results in a temperature sensitive repressor protein; at lower temperatures at 37°C and below, λ prophages with the *c*I*857* allele form stable lysogens, and switch to lytic growth at temperatures above 37°C (Sussman & Jacob, 1962). This occurs without any requirement for DNA damage or an SOS response. The *c*I *ind1* allele (Jacob & Campbell, 1959) results in an E117K amino acid change in the CI repressor protein (Gimble & Sauer, 1985) that prevents SOS mediated RecA* coprotease autocleavage of the CI repressor, making the prophage uninducible by DNA damage.

Baek et al. (2003) noticed that a λ lysogen with the *rexA* and *rexB* genes replaced with *gfp* was less inducible at low ultraviolet (UV) doses than a lysogen with *rexA* and *rexB* intact, suggesting that one or both of the Rex proteins have a role in prophage induction. We have further explored the effects of RexA and/or RexB on repression and the induction of λ from the repressed prophage state. Our findings suggest that RexA stabilizes the nonimmune state. RexB appears to antagonize the effect of RexA and may be involved in establishment of the phage DNA replication complex once the transition to lytic growth occurs. Results showing that the Rex system may assist in the establishment of phage replication are found in Supporting Information Results and Discussion and Table S1.

## Results

### RexA and RexB proteins modulate the level of phage yield in response to UV induction

We compared the ultraviolet inducibility of λ*c*I^+^ lysogens, defective in RexA and/or RexB functions, to each other and to wild type Rex+ lysogens (Figure 2A). The lysogen defective for RexA function, LT1676, retains the *P*_LIT_ promoter and expresses RexB fully from both *P*_RM_ and *P*_LIT_ (see bacterial strains in Table 1). Relative to a wild type λ *rexA^+^ rexB^+^* lysogen (filled circles), the three lysogens defective for Rex function induce less well at low UV doses, but all induce like wild type at the highest doses (Figure 2A). λ *c*I^+^ lysogens defective for RexA function are very defective for induction at low levels of irradiation irrespective of whether the phage expresses RexB function. In fact, the RexA mutant (open triangles) is as impaired for induction at low UV doses as a lysogen lacking both RexA and RexB functions (open circles). In contrast, the lysogen defective for only RexB function (filled triangles) is similar to wild type in responding to low levels of UV and shows only a modest reduction in levels of phage produced, i.e., pfu/ml in Figure 2A. In summary, these results demonstrate a positive activity of RexA in promoting induction of the lysogen when switching to the lytic state in response to low levels of DNA damage. There is a small positive effect of RexB when it is co-expressed with RexA. All four lysogens, independent of their *rex* genotype, respond to the same extent to a fully inducing dose of UV (~15 J/m^2^) (see Figure 2A). Researchers other than Baek *et al*. (2003) may have missed this effect of Rex functions on UV induction since the historic dose used to induce λ has been ~15 J/m^2^ (Coetzee & Pollard, 1974). We also find that expression of RexA and RexB from the arabinose operon can stimulate UV induction of a λ*cI*^+^ *rexA*^−^ *rexB*^−^ lysogen (Figure 2B, open diamonds), confirming that the λ Rex functions can complement *in trans* to alter the basic behavior of the CI/Cro bistable switch when responding to lower levels of UV.

**Figure 2.**
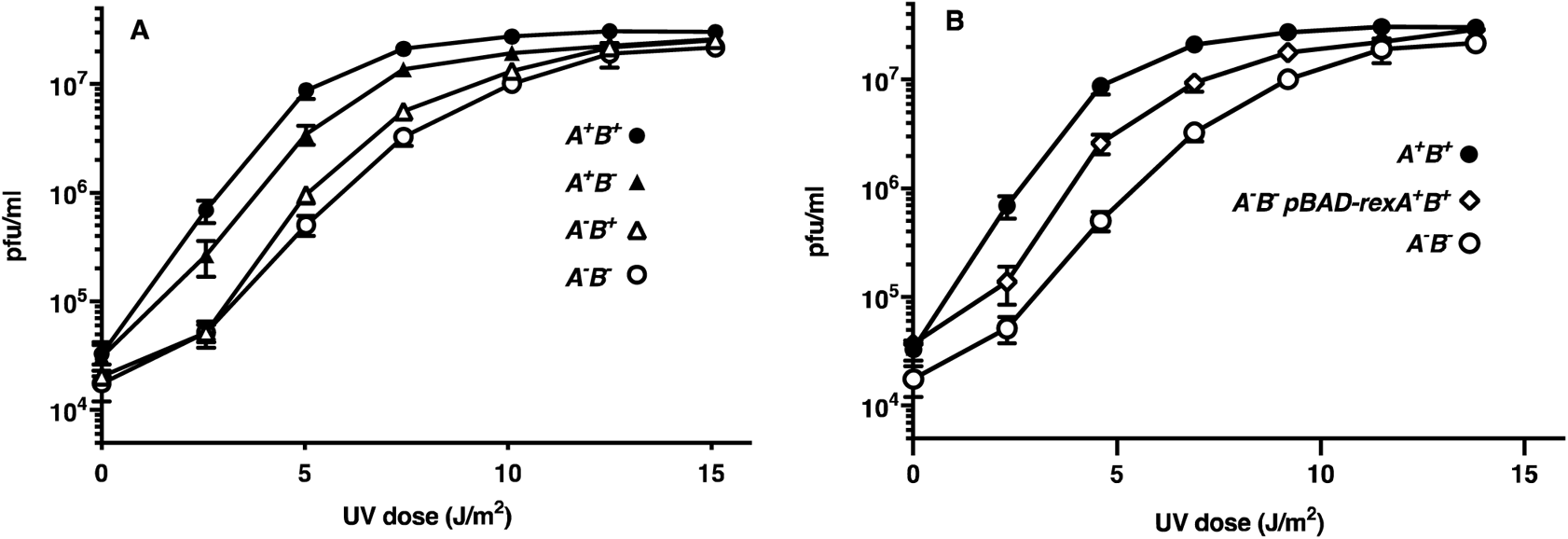
RexA^+^ enhances UV induction of a λ *c*I^+^ prophage at low UV doses. UV induction of λ lysogens was performed as described in Experimental Procedures. **A.** LT445 is MG1655(λ *c*I^+^ *rexA*^+^ *rexB*^+^) (●) (n=7); LT1676 is MG1655(λ*c*I^+^ *rexA<>cat rexB*^+^) (Δ) (n=3); LT1677 is MG1655(λ*c*I^+^ *rexA*^+^ *rexB<>cat*) (▴) (n=6), LT1678 is MG1655(λ*c*I^+^ *rexA*-*rexB<>cat*) (○) (n=3). Titers of the lysates on strain A584 were determined at t=0 and the time points indicated following UV irradiation. **B.** LT445 is MG1655(λ) (●); LT1678 is MG1655(λ *rexA*-*rexB<>cat*) (○) (n=3); LT2109 is MG1655(λ *rexA*-*rexB<>cat*)/*P*_BAD_-*rexA^+^ rexB^+^* (□) (n=4). Data have been analyzed by the standard error of the mean (s.e.m). The number of trials is indicated by (n).

**Table 1.** *Escherichia coli* K-12 Strains.

| Strain | Relevant Genotype | Reference/construction |
| --- | --- | --- |
| A584 | YMC [*trp::Tn10 tyrT (supF)*] | D. Ennis via F. W. Stahl |
| C600 | *tonA21 thi-1 thr-1 leuB6 lacY1 rfbC1 fhuA1 glnV44 (supE)* | D. Court laboratory |
| LT445 | MG1655($\lambda$ *cI<sup>+</sup> rexB<sup>+</sup> rexA<sup>+</sup>*) monolysogen | Thomason et al., 2019 |
| LT1055 | MG1655 Δ*lacI-kan luc-N pLoL rexB<sup>+</sup> rexA<sup>+</sup> cI857ind1 pRoR cro27 cII-lacZYA* | this work |
| LT1395 | MG1655 Δ*lacI-kan luc-N pLoL rexB<sup>+</sup> rexA<>cat cI857ind1 pRoR cro27 cII-lacZYA* | this work |
| LT1657 | MG1655 Δ*lacI-kan luc-N pLoL rexB<sup>+</sup> rexA<sup>+</sup> cI<sup>+</sup> pRoR cro<sup>+</sup> cII-lacZYA* | |
| LT1659 | MG1655 Δ*lacI-kan luc-N pLoL rexB<sup>+</sup> rexA<>cat cI<sup>+</sup> pRoR cro<sup>+</sup> cII-lacZYA* | |
| LT1676 | MG1655($\lambda$ *cI<sup>+</sup> ind<sup>+</sup> rexB<sup>+</sup> rexA<>cat*) monolysogen | Thomason et al., 2019 |
| LT1677 | MG1655($\lambda$ *cI<sup>+</sup> ind<sup>+</sup> rexB<>cat rexA<sup>+</sup>*) monolysogen | Thomason et al., 2019 |
| LT1678 | MG1655($\lambda$ *cI<sup>+</sup> ind<sup>+</sup> (rexAB)<>cat*) monolysogen | Thomason et al., 2019 |
| LT1684 | MG1655($\lambda$ cI857 *ind1* *rexB*<sup>+</sup>*rexA*<sup>+</sup>)<br>*lamB*<>*kan* monolysogen | this work |
| LT1865 | MG1655 Δ*lacI-kan luc-N pLoL*<br>*rexB*<>*cat rexA*<sup>+</sup> cI857*ind1 pRoR cro27*<br>*cII-lacZYA* | this work |
| LT1866 | MG1655 Δ*lacI-kan luc-N pLoL* (*rexB*-<br>*rexA*<>*cat*) cI857*ind1 pRoR cro27 cII-*<br>*lacZYA* | this work |
| LT1886 | MG1655 Δ*lacI-kan luc-N pLoL rexB*<sup>+</sup><br>*rexA*<sup>+</sup> cI857 *pRoR cro*<sup>+</sup> *cII-lacZYA* | this work |
| LT1887 | MG1655 Δ*lacI-kan luc-N pLoL rexB*<sup>+</sup><br>*rexA*<>*cat cI857ind1 pRoR cro*<sup>+</sup> *cII-*<br>*lacZYA* | this work |
| LT1891 | MG1655 Δ*lacI-kan luc-N pLoL*<br>*rexB*<>*cat rexA*<sup>+</sup> cI857*ind1 pRoR cro*<sup>+</sup><br>*cII-lacZYA* | this work |
| LT1892 | MG1655 Δ*lacI-kan luc-N pLoL* (*rexB*-<br>*rexA*<>*cat*) cI857*ind1 pRoR cro*<sup>+</sup> *cII-*<br>*lacZYA* | this work |
| LT1895 | MG1655 Δ*lacI-kan luc-N pLoL*<br>*rexB*<>*cat rexA*<sup>+</sup> cI<sup>+</sup> *pRoR cro*<sup>+</sup> *cII-*<br>*lacZYA* | |
| LT1897 | MG1655 Δ*lacI-kan luc-N pLoL* (*rexB*-<br>*rexA*<>*cat*) cI<sup>+</sup> *pRoR cro*<sup>+</sup> *cII-lacZYA* | |
| LT2063 | LT1886 $\Delta$ (*srlA- recA*)::Tn10 | this work |
| LT2064 | LT1887 $\Delta$ (*srlA*- *recA*)::*Tn10* | this work |
| LT2065 | LT1891 $\Delta$ (*srlA*- *recA*)::*Tn10* | this work |
| LT2066 | LT1892 $\Delta$ (*srlA*- *recA*)::*Tn10* | this work |
| LT2109 | LT1678 *P*<sub>BAD</sub>-(*rexA*<sup>+</sup> *rexB*)<sup>+</sup> | this work |
| LT2319 | MG1655($\lambda$ *cI857 ind1 rexB*<sup>+</sup>*rexA*<>*cat*)<br>*lamB*<>*kan* monolysogen | this work |
| LT2320 | MG1655($\lambda$ *cI857 ind1 rexB*<>*cat rexA*<sup>+</sup>)<br>*lamB*<>*kan* monolysogen | this work |
| LT2321 | MG1655($\lambda$ *cI857 ind1(rexB rexA)*<>*cat*)<br>*lamB*<>*kan* monolysogen | this work |

### RexA enhances the spontaneous transition to the nonimmune state, as monitored by a PR lacZ reporter

Our data show that the RexA protein stimulates lytic growth and phage production at suboptimal UV doses. We next asked whether the positive effect of RexA on prophage induction requires SOS-induced autocleavage of CI protein by determining whether RexA stimulates when CI repressor is not cleavable, i.e., contains the *ind1* mutation, or in RecA-mutant cells defective for the SOS response. We used the *c*I*857* temperature sensitive repressor allele for these experiments, since Toothman and Herskowitz (1980) previously noticed enhanced Rex-dependent effects in a *c*I*857* repressor background at temperatures where CI-mediated repression is active. Reichardt (Reichardt, 1975) showed that repressor protein levels are higher in *c*I*857* lysogens than in *c*I^+^ lysogens. This higher level of *c*I*857* protein is made in response to the autoregulatory properties of the *P*_RM_ promoter in order to maintain a constant level of CI repression. Thus, more *P*_RM_ activated transcription will also generate more RexA/B protein and would account for enhanced RexA activity.

*E. coli* strains containing λ *c*I*857 ind1* constructs with either the cro+ or cro^−^ alleles were modified to carry the four combinations of the rexA and rexB genes described in the legend of Figure 2A. These eight constructs contain only the phage immunity region as shown in Figure 3 and are nonlethal; such constructs have been described by Svenningsen et al. (2005). Cultures of the Cro^+^ strains were grown in L broth overnight, diluted and spread for individual colonies on MacConkey Lactose agar. The indicator plates were incubated at low temperature where CI857 repressor is active. After 2-3 days, red papillae formed (Figure 4) in the bacterial colonies, which indicated that within the colony there was a clonal population of cells expressing β-galactosidase from *P*_R_-*c*II-*lacZ* transcripts (see Figure 3). These results suggested that either mutations were occurring within the *c*I repressor gene allowing *P_R_* transcription of *lacZ*, or that an epigenetic change had occurred to lift CI repression and allow Cro repression (Ptashne et al., 1980), i.e., the bistable switch had flipped from the immune state to the lytic state in these papillae. We found that colonies from the two strains having RexA function (LT1886 and LT1891) underwent more papillation than the two strains defective for RexA (LT1887 and LT1892) (Figure 4A), in experiments controlled for temperature and time. Since the non-cleavable *c*I *ind1*-allele is present in this set of four strains, CI repressor autocleavage is not required for this Rex phenotype. Congruent with this observation, RexA-dependent patterns of papillation occur whether or not the strains are functional for RecA (compare Figure 4A vs 4B). Thus, RexA modulation of the bistable switch does not act through the SOS pathway of CI repressor autocleavage. Confirmation that the rare papillae arise from epigenetic changes rather than from null mutations in the *c*I repressor gene is presented below.

**Figure 3.**
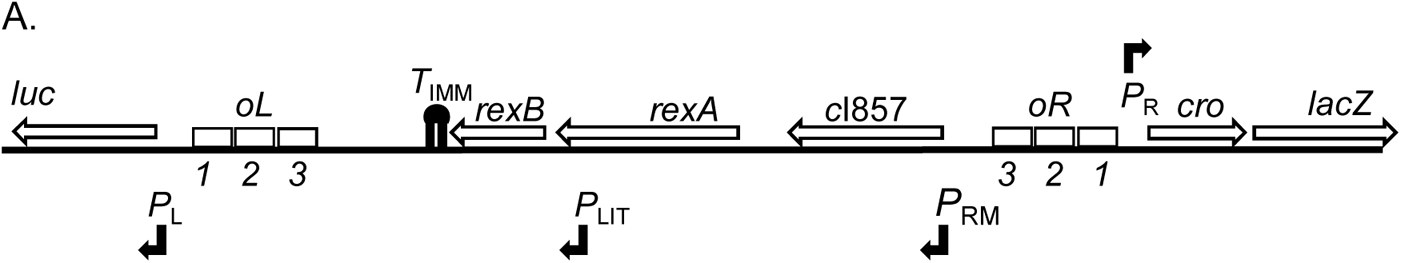
Genetic map of dual *P*_L_ *P*_R_ reporter. The phage λ immunity region has been inserted within the *E. coli lac* operon such that expression of *lacZ* is driven from the *P*_R_ lytic promoter (Svenningsen 2005) with the *lacI* gene and the *lac* promoter being replaced by *P*_R_. This leaves the *lacZ* ribosome-binding site and the rest of the *lacZYA* operon intact. Versions of this reporter with the temperature sensitive *c*I*857* repressor were used to determine the effects of RexA and/or RexB functions on induction and the switch to activate *P*_L_ and *P*_R_ promoter transcription. Single colonies were grown on MacConkey Lactose indicator medium. When the switch is in the CI-repressed or immune state, colonies do not express LacZ and are white. When the switch is in the Cro-repressed nonimmune state, *lacZ* is expressed from *P*_R_ and the colonies are red. The firefly luciferase gene *luc* replaces the λ *N* gene beyond the *P*_L_ promoter.

**Figure 4.**
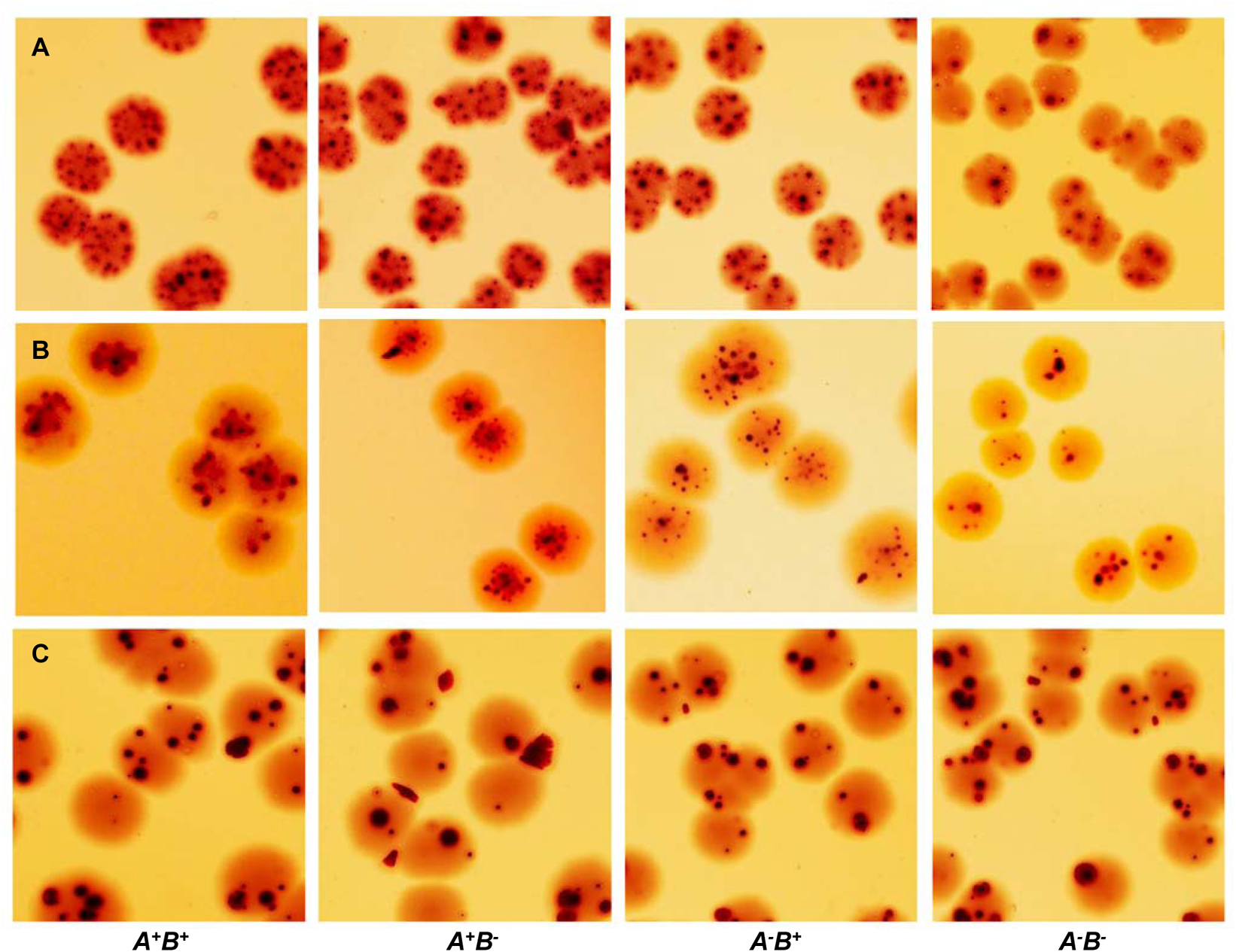
RexA promotes the transition to nonimmune state within individual colonies. In these pictures, colonies of strains containing the *P*_L_ and *P*_R_ reporters with the temperature sensitive *c*I*857* repressor allele were plated on MacConkey Lactose agar and incubated at 32°-34°C. All colonies are white after one day of incubation but develop red papillae after two days, indicative of a transition to the lytic state by cells within the colony and consequent expression of *lacZ* from *P*_R_. The *rex* genotypes are indicated, and the strain numbers are shown below. **A.** Top row: the strains (LT1886, LT1887, LT1891, and LT1892) display different papillation levels dependent on the genotype of the *rexA* and *rexB* genes. **B.** Middle row: The *recA* mutant strains (LT2063, LT2064, LT2065, and LT2066) also display variable papillation based on *rex* genotype. **C.** Bottom row: The *cro27* mutant strains of λ (LT1055, LT1395, LT1865, and LT1866) all papillate similarly, regardless of *rex* genotype.

### A functional Cro gene is required for Rex-dependent papillation

The Cro repressor is an integral component of the bistable switch and is required for lytic growth (Folkmanis et al., 1977; Lee, Lewis, & Adhya, 2018). We found that the Rex-dependent effects on colony papillation are only observed when Cro function is present. When the RecA^+^ reporter strains with the four combinations of *rexA* and *rexB* are defective for the Cro repressor, the papillation phenotype becomes independent of Rex function, with papillae occurring at a lower frequency than in the Cro^+^ strains (Figure 4C). Sequence analysis of four red papillae from each of the Cro mutant strains revealed the presence of mutations in the *c*I gene which render them unable to maintain the immune state. In contrast, no *c*I mutations were found under RecA^+^ Cro^+^ conditions when four red papillae of each genotype were sequenced.

### RexA promotes stability of the nonimmune state

Twenty different papillae from each of the eight strains shown in Figure 4A and 4B were struck on MacConkey Lactose agar for purification. When each of these purified red colonies is grown in L broth and then spread on MacConkey Lactose agar, rare white colonies appear among the red colonies on the plate. Thus, not only do immune (white) colonies contain papillae that have switched to the nonimmune (red) state, but once switched, they can also switch back to the immune (white) state. The percentage of switched white colonies within each culture was determined for each genotype (Figure 5). The most stable nonimmune state is in the cells with wild type RexA and RexB functions present. This is true whether cells are *recA*^+^ or *recA*^−^, with 0.9% of *recA*^+^ (Figure 5A) and 1.7% of *recA*^−^ (Figure 5B) cells switching to the immune state. RexA function exerts a strong positive effect to maintain the nonimmune state even in the absence of RexB, with 1.6% of *recA*^+^ (Figure 5A) and 2.7% of *recA*^−^ (Figure 5B) cells in each culture reverting to the immune state. Cells lacking RexA function display higher rates of reversion to the immune state whether or not RexB is present and whether or not host RecA protein is present. We observed small differences due to removal of host RecA protein (see explanation in Fig S1 of Supporting Information), but our main observation, that the presence of RexA enhances stability of the lytic state of the bistable switch, is independent of RecA function.

**Figure 5.**
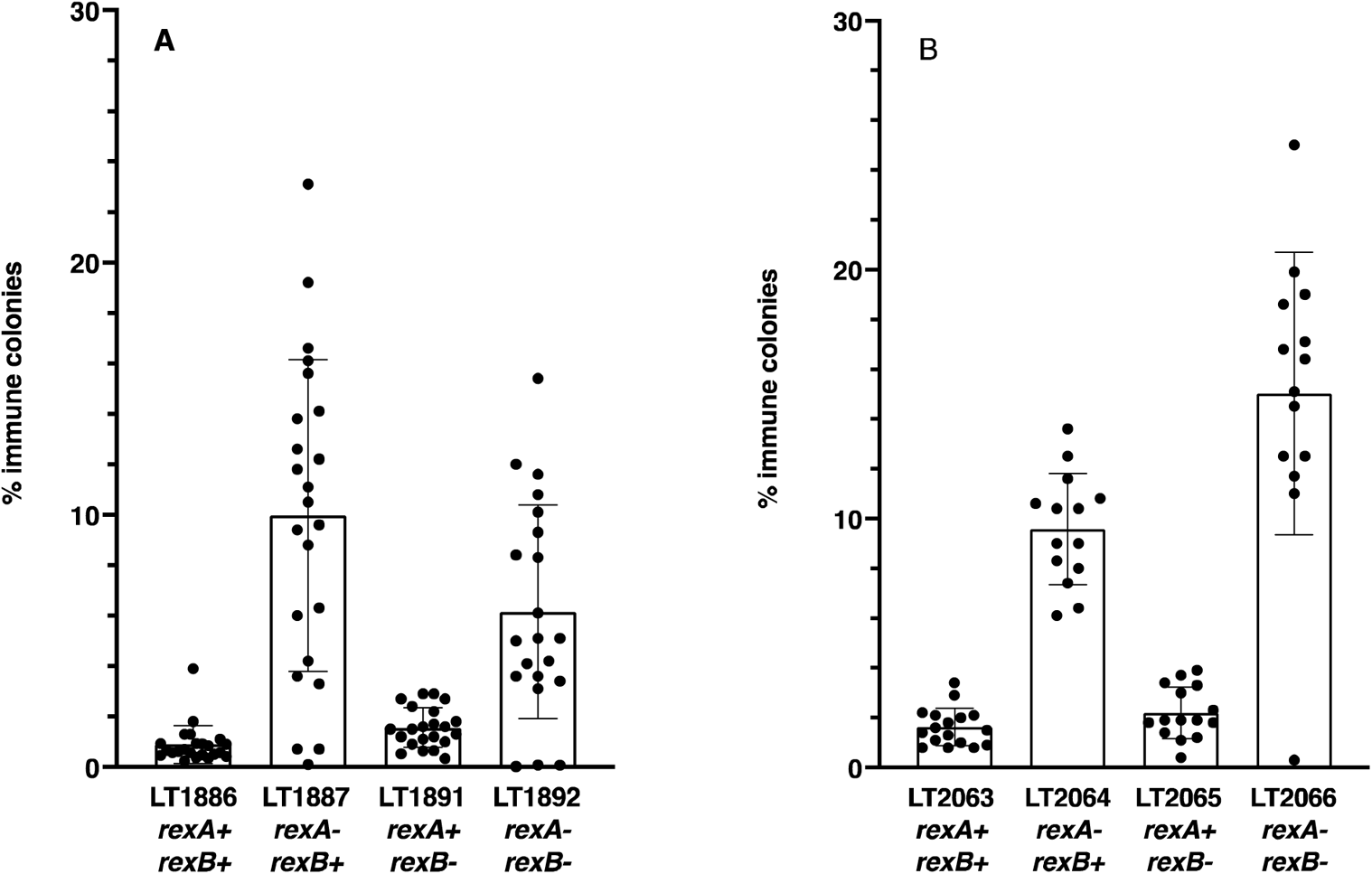
RexA function stabilizes the non-immune state. RecA^+^ and RecA^−^ colonies containing the Cro^+^ *P*_R_ reporter pictured in Fig. 3 were analyzed to determine the effect of RexA and RexB on the frequency of returning from the non-immune state to the immune state in the presence of Cro repression. The y-axis indicates the percentage of immune (white) colonies arising during overnight growth of a culture that initially carried a reporter in the non-immune (red) state. The final number of colonies analyzed for each genotype is indicated. The error bars show the standard deviation (s.d.). **A.** RecA^+^ strains: LT1886, *rexA^+^ rexB^+^*(n=18); LT1887, *rexA^−^ rexB^+^* (n=21); LT1891, *rexA^+^ rexB^−^* (n=22); LT1892, *rexA^−^ rexB^−^* (n=20). **B.** Δ*recA* strains: LT2063, *rexA^+^ rexB^+^* (n=17); LT2064, *rexA^−^ rexB^+^* (n=14); LT2065, *rexA^+^ rexB^−^* (n=15); LT2066, *rexA^−^ rexB^−^* (n=14).

We carried out a similar analysis with Lac^+^ red papillae arising in the RecA^+^ *cro27* mutant strains LT1055, LT1395, LT1865, and LT1866, which contain the four combinations of RexA and RexB (see Figure 4C) and are defective for Cro function (Eisen, Brachet, Pereira da Silva, & Jacob, 1970). When the red colonies are grown in L broth and dilutions plated on MacConkey Lactose agar, no white colonies were found among thousands of red colonies. This indicates that the *cro27* Lac^+^ colonies contain mutations in *c*I, which inactivate repressor function as discussed earlier. Our data indicate that red papillae due to *c*I mutations do occur but compose a small minority of the total papillae under Cro^+^ conditions, whereas, under Cro^−^ conditions, we found only *c*I mutations.

### RexA^+^ strains show earlier expression of the *P_L_* lytic promoter during the transition to the nonimmune state

We looked for Rex-dependent transcription differences in expression of a reporter gene under control of the lytic promoter *P*_L_ during transition of the bistable switch to the lytic state. Strains LT1657, LT1659, LT1895 and LT1897 are similar to those used for the papillation experiment (Figure 4A) but contain a wildtype CI repressor (*c*I^+^*ind^+^*); thus, in these strains, the nonimmune state is inducible by DNA damage but not by high temperature.

A low dose of Mitomycin C (MC) was added to log phase cultures of these strains to cause DNA damage and trigger RecA*-dependent autocleavage of CI repressor and activation of *P*_L_- and *P*_R_-mediated transcription. Once induced by MC, Cro expression from *P*_R_ results in repression of *P*_RM_, completing the switch from the lysogenic to the lytic state. The results are shown in Figure 6.

**Figure 6.**
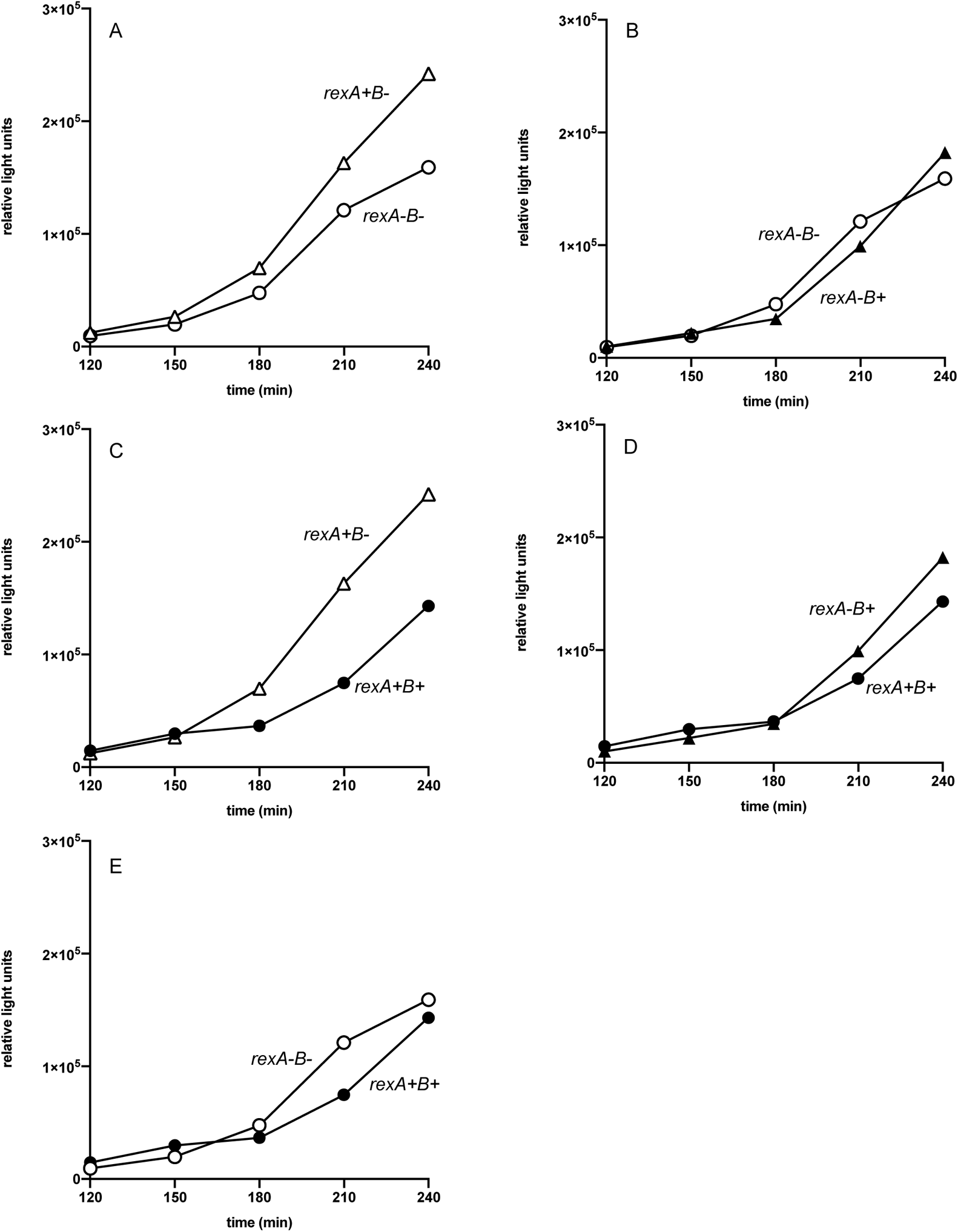
Monitoring luciferase activity from *P*_L_ *N-luc* after inducing DNA damage. After addition of Mitomycin C, expression of firefly luciferase from the *P*_L_ promoter (*P*_L_ *N-luc*) was monitored over time for four strains carrying the lambda immunity region and different *rexA* and *rexB* mutations. Relative light units are shown on the y-axis and time in minutes on the x-axis. The DNA damage inducer, Mitomycin C, was added at t=0. For the first ~two hours all points are congruent, but Rex-dependent genotypic differences arise in luciferase expression are apparent after ~150 min. (A). LT1895, *rexA^+^ rexB<>cat* (Δ) vs. LT1897, (*rexA rexB*)*<>cat* (○). (B). LT1659, *rexA<>cat rexB^+^* (▴) vs. LT1897, (*rexA rexB*)*<>cat* (○).(C). LT1895, *rexA^+^ rexB<>cat* (Δ) vs. LT1657, *rexA^+^ rexB^+^* (●). (D). LT1659, *rexA<>cat rexB^+^* (▴) vs. LT1657, *rexA^+^ rexB^+^* (●). (E). LT1657, *rexA^+^ rexB^+^* (●) vs. LT1897, (*rexA rexB*)*<>cat* (○). The experiment was repeated three times with duplicate technical replicates each time; one representative experiment is shown.

The effect of RexA and/or RexB function on expression of luciferase from the lytic promoter *P*_L_ was monitored over time in the four different strains. The CI and Cro repressors were wild type in all cases. When RexA function is present and RexB is absent (Figure 6A, strain LT1895, open triangles), luciferase expression from the lytic promoter *P*_L_ occurs reproducibly earlier than when both RexA and RexB are absent (strain LT1897, open circles); this again demonstrates the positive effect of RexA on the transition to the lytic state. In contrast, when RexB function is present and RexA is absent (Figure 6B, strain LT1659, closed triangles), the kinetics of luciferase expression from *P*_L_ are similar to those observed when both RexA and RexB are absent. Thus, expression of RexB alone does not alter the timing of *P*_L_ activation. However, when both RexA and RexB proteins are present (Figure 6C, strain LT1657, closed circles), *P*_L_ expression is delayed relative to the strain with only RexA function (open triangles). Thus, when present, RexB can inhibit RexA stimulation of luciferase from *P*_L_. This experiment illustrates the independent and opposing effects that RexA and RexB have on the transition to the lytic state by affecting the timing of expression of the lytic *P*_L_ promoter. Note that the earliest time points show no genotypic differences, suggesting that RexA does not act until DNA damage has occurred, CI repressor levels are reduced, and prophage induction has begun.

### Rex effects on spontaneous phage release of *c*I857 *ind1* lysogens

We have shown that RexA function stimulates switching from the immune to the nonimmune state when Cro function is present by monitoring the expression of reporter genes from the major rightward (*P*_R_) and leftward (*P*_L_) lytic promoters in a defective prophage. We next looked for effects of Rex function on the bistable immunity switch in λ lysogens carrying a complete prophage. λ lysogens, when grown in culture, are relatively stable and rarely release phage. There is, however, a low level of phage release. Some of the phage released are rare *c*I mutants that have simply lost repressor activity, but the vast majority are genetically identical to the original prophage and arise from a cell in which λ had switched from the immune to the non-immune state; the two types are distinguishable by plaque morphology (Little & Michalowski, 2010). We tested the effects of RexA and/or RexB on this spontaneous phage release using *c*I*857* lysogens with the *ind1* mutation to block SOS-mediated prophage induction (see Figure 7).

**Figure 7.**
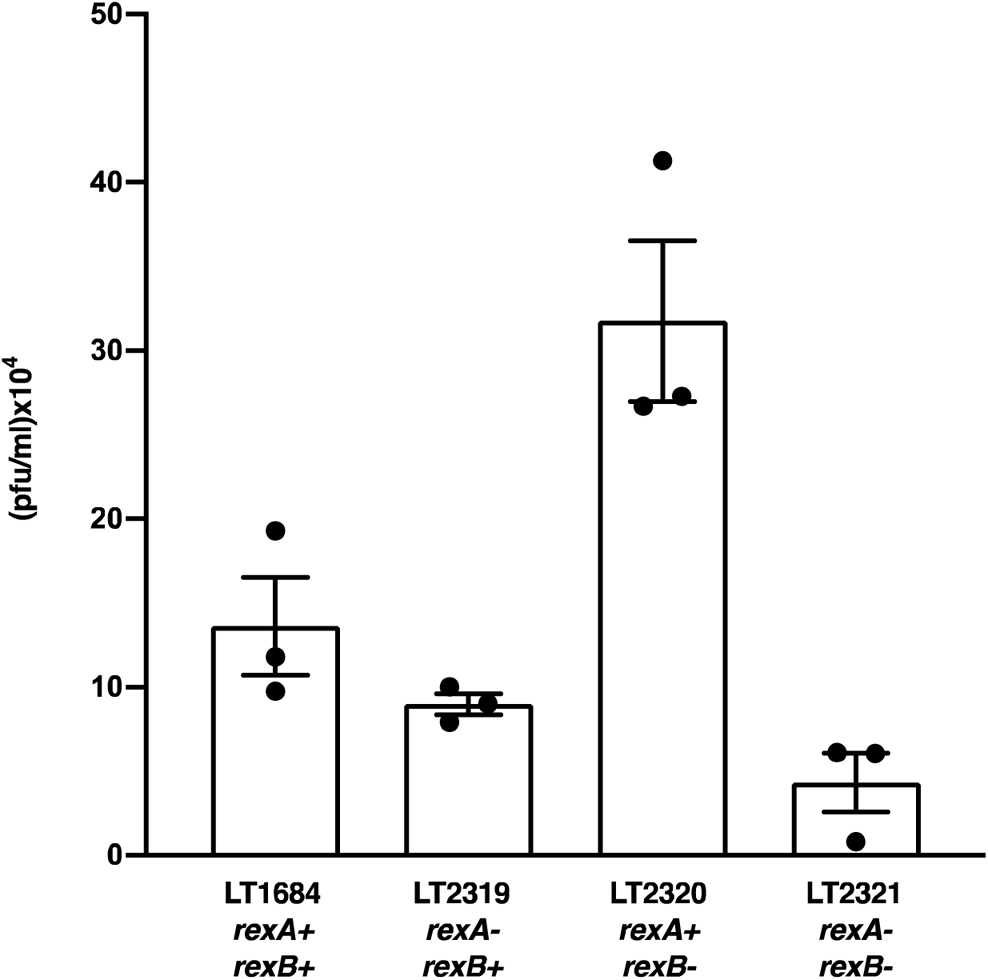
RexA and RexB affect the level of spontaneous phage release from λ *c*I*857 ind1* lysogens. The bar graph shows the plaque-forming units (PFU) per ml of spontaneously released phage particles arising during 32°C growth of lysogenic cultures. Strain numbers and *rex* genotypes for each culture are indicated below the bars. Three independent repetitions of each experiment were performed; error bars represent the standard deviation (s.d.).

We find that RexA enhances the transition to lytic growth. This RexA effect is most evident in the absence of RexB function (Figure 7, compare LT2320 and LT2321), where removal of RexA activity by mutation results in a >7-fold reduction in free phage titer. As observed previously, this RexA effect is nearly absent when RexB function is present: compare the *rexA^+^* LT1684 with the *rexA* mutant LT2319; the levels of free phage in these two lysogens varies by <2-fold (see Figure 7). Clear plaques result from *c*I mutations that prevent lysogeny and account for less than 5% of the total phage yield for all genotypes. Thus, while demonstrating the positive effect of RexA protein on the transition to lytic growth, the data also reveal that RexB protein antagonizes RexA function.

### Two-hybrid analysis of protein-protein interactions between RexA, RexB and the CI and Cro repressors

Protein-protein interactions occurring between RexA, RexB, and the phage repressors CI and Cro might affect the operation of the bistable switch. Thus, we looked for such interactions using the Bacterial Adenylate Cyclase Two Hybrid (BACTH) system (Karimova, Gauliard, Davi, Ouellette, & Ladant, 2017; Ouellette, Karimova, Davi, & Ladant, 2017). This two-hybrid analysis is performed in an *E. coli cya* mutant lacking adenylate cyclase function, using compatible plasmids expressing the T25 and T18 adenylate cyclase domains from *Bordatella pertussis* fused to either the N- or C-terminus of proteins of interest. Interaction between the two proteins being tested results in association of the two cyclase domains, giving cyclase activity and consequent cAMP production. We previously used this system to show that RexA and RexB interact with themselves and each other (Thomason et al., 2019). For that study we generated eight plasmids that express the *rexA* and *rexB* genes fused in frame at either the N- or C-terminus to each of two different cyclase domains. Here we made eight additional plasmids by inserting the phage repressor genes, *c*I and *cro*, into the four cyclase vectors in the same manner. We then introduced plasmid pairs to be tested for interaction into the *cya* mutant host, BTH101 (Table S2 A-E). To ensure that only meaningful interactions were included in the analysis, we first screened strains with the pairs of plasmids for the ability to confer growth on minimal maltose (see pictures accompanying Table S2 A-E), since activation of the *mal* genes depends strongly on cAMP (Raibaud, Vidal-Ingigliardi, & Kolb, 1991). We then measured β-galactosidase production from the bacterial *lacZ* gene for each of these positive plasmid pairs, since expression of the *lac* operon also requires cAMP for promoter activity (see Table 2). The results confirm that both RexA and RexB proteins interact with the CI repressor. For RexA and CI, only proteins with N-terminal cyclase tags allowed growth on maltose, suggesting that the C-terminal domains of these two proteins interact. It is known that CI repressor dimers form via an interaction domain located in the repressor C-terminus (Beckett, Burz, Ackers, & Sauer, 1993; Burz & Ackers, 1994). Thus, it is not surprising that these CI hybrid proteins form dimers with the cyclase tag on the N-terminus. Robust β-galactosidase values suggest that CI repressor with an N-terminal cyclase tag also interacts well with RexB protein. The cyclase tags can be at either end of RexB, which is an inner membrane protein, and predicted to have both N- and C-termini in the cytoplasm (Parma et al., 1992). β-galactosidase assays testing interaction of the Cro repressor with RexA and RexB proteins (Table 2) show about half the level of signal that we found for CI repressor-RexA/B interaction.

**Table 2.**
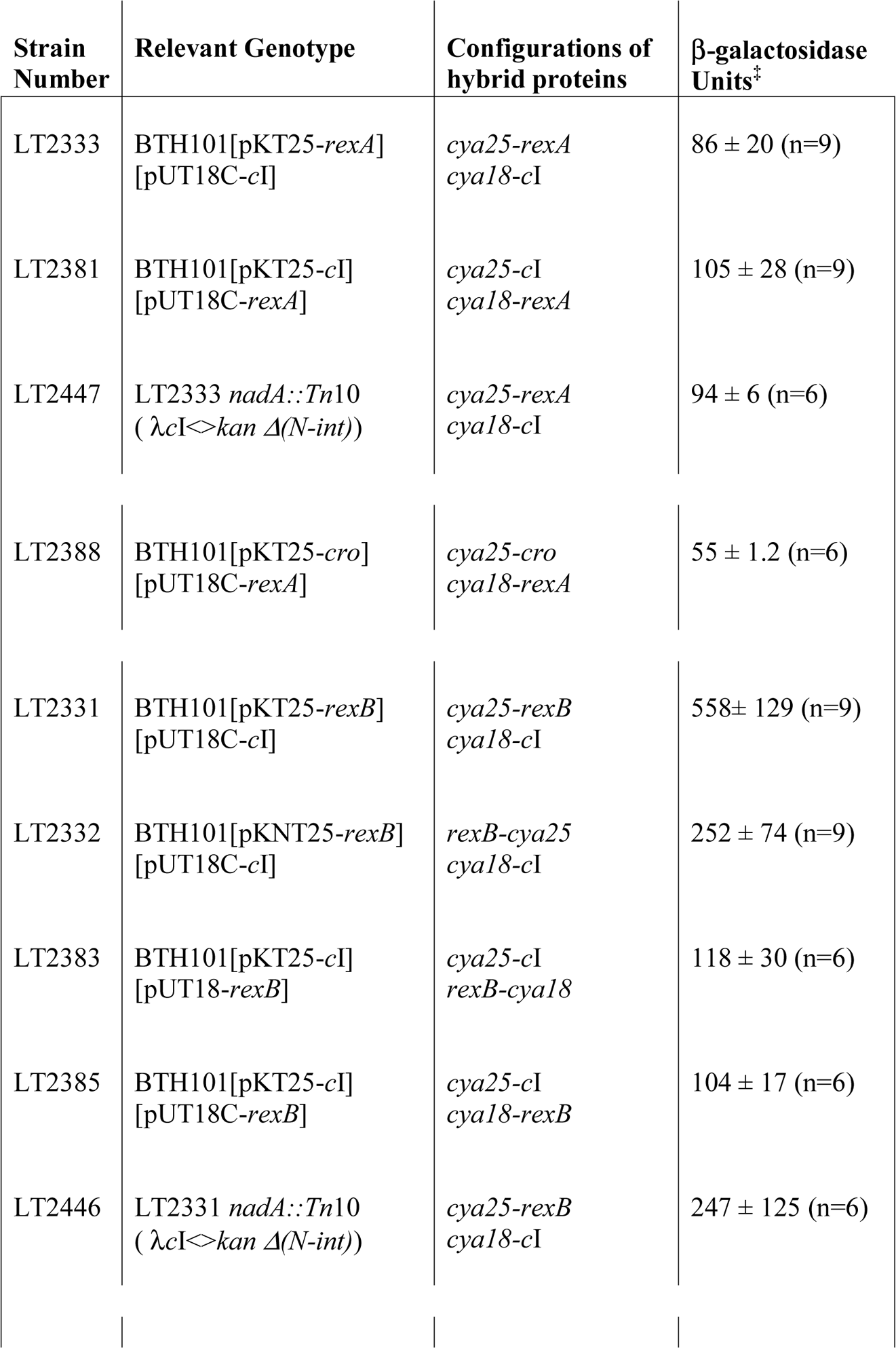
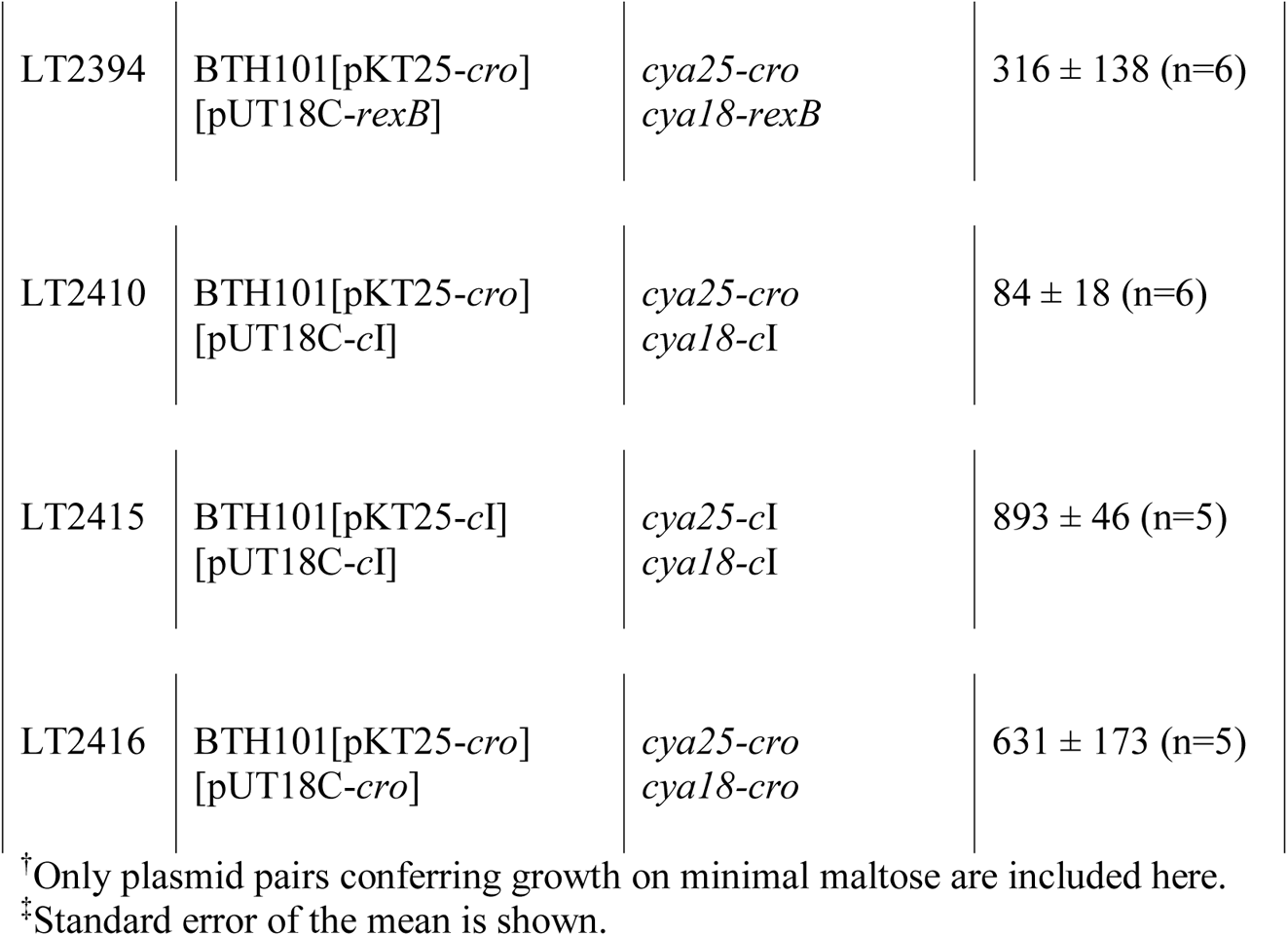
β-galactosidase measurements for BACTH system demonstrates protein-protein interaction between RexA and RexB with CI and Cro repressors^†^.

The plasmids expressing the hybrid CI repressor confer immunity to λ, demonstrating that the CI fusion proteins are able to bind to the operator sites at the lytic promoters. Since binding to the operators might change protein-protein interactions and thus the β-galactosidase results, we tested a subset of the plasmid pairs in a derivative of the *cya* mutant strain having the phage operator sites present on the bacterial chromosome. However, we saw no significant differences in β-galactosidase levels with or without the repressor operator sites present (see Table 2, LT2333 vs LT2446 and LT2331 vs LT2447).

### RexA is a non-specific DNA binding protein that forms stable complexes with CI oligomers *in vitro*

Our BACTH data demonstrate that RexA and RexB can associate with CI *in vivo*. To study these interactions directly, we sought to purify RexA and RexB recombinantly from *E. coli* and test whether they could bind CI *in vitro*. RexA can be purified in milligram quantities as a soluble, monodispersed dimer (Figure 8A). We observe no evidence of subunit exchange in solution, suggesting that dimerization occurs during protein folding and the dimer remains stably associated. All attempts to purify RexB were unsuccessful, regardless of the tag used or how the protein was expressed.

**Figure 8.**
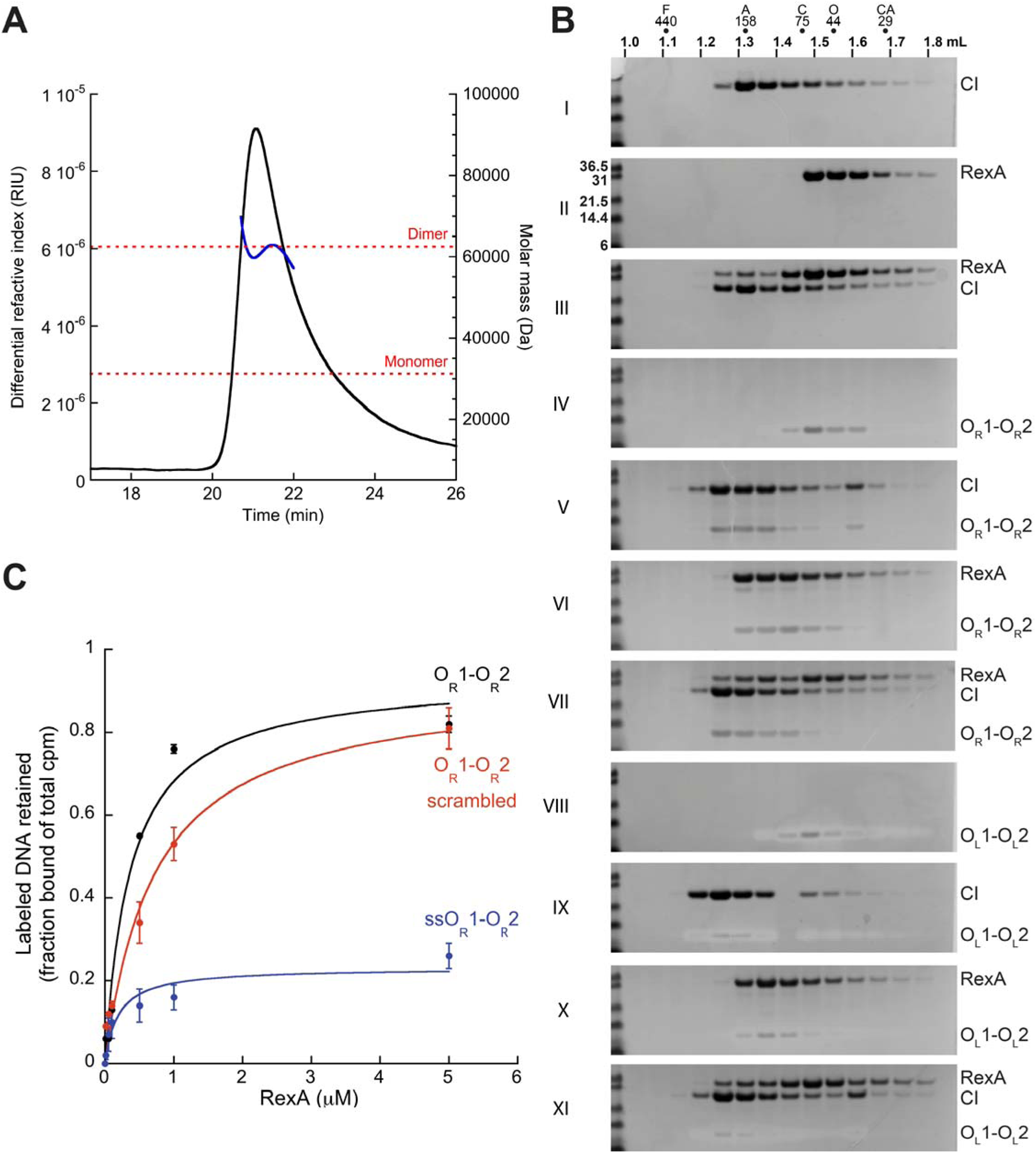
RexA forms stable complexes with CI and DNA *in vitro*. **A.** SEC-MALS analysis of purified RexA. UV trace (black) and measured mass based on light scattering (blue) are shown. **B.** SEC analysis of RexA and CI protein-protein and protein-DNA interactions. SDS-PAGE gels (silver-stained for DNA and Coomassie-stained for protein) from individual SEC injections are numbered with Roman numerals and shown to visualize shifts in retention volume off of SEC in response to different conditions. Molecular weight standards in kDa are shown in the first lane of each gel with samples labeled on the right. All samples were run on a Superdex 200 PC 3.2 column (GE). The elution volume across the fractions is marked above along with the relative positions of molecular weight standards (F, ferritin, 440 kDa; A, aldolase, 158 kDa; C, conalbumin, 75 kDa; O, ovalbumin, 44 kDa; CA, carbonic anhydrase, 29 kDa). A leftward shift of the bands indicates formation of a larger molecular weight species and is associated with complex formation. See Materials and Methods and Table S3 for DNA substrate preparation and oligonucleotide sequences, respectively. **C.** Filter binding analysis of RexA interactions with different DNA substrates. Substrate nomenclature: OR1-OR2, double-stranded DNA containing wildtype O_R_1 and O_R_2 operator sites; ss, single-stranded DNA; scrambled, mutated substrate altering operator site sequences. Binding was performed with wildtype RexA at 30°C for 10 min in a 30 μL reaction mixture containing 14.5 nM unlabeled DNA and 0.5 nM labelled DNA. Samples were filtered through KOH-treated nitrocellulose and binding was assessed by scintillation counting. The data points represent the averages of at least three independent experiments (mean ± standard deviation) and were compared to a negative control to determine fraction bound.

Analytical size exclusion chromatography (SEC) shows that purified CI repressor elutes earlier than the RexA dimer under the same experimental conditions (Figure 8B, gels I and II), consistent with the ability of CI to form larger oligomers (Bell, Frescura, Hochschild, & Lewis, 2000; Dodd, Perkins, Tsemitsidis, & Egan, 2001; Révet, von Wilcken-Bergmann, Bessert, Barker, & Müller-Hill, 1999; Stayrook, Jaru-Ampornpan, Ni, Hochschild, & Lewis, 2008). When the assembled CI multimers are mixed with RexA dimers, a portion of RexA co-elutes with CI and both proteins are enriched in the heaviest, early fractions (8B, gel III versus I and II), indicating that RexA can form stable complexes with CI in solution.

CI also forms complexes with double-stranded DNA (dsDNA) containing either the λ *O*_L_ or *O*_R_ operator sequences (Figure 8B, gels I, IV, and V and gels I, VIII, and IX), as evidenced by the leftward shift of the DNA bands when CI is added (Maniatis & Ptashne, 1973). Surprisingly, RexA also shifts to a larger species in the presence of these substrates (Figure 8B, gels II, IV, and VI and gels II, VIII, and X; Supporting Information Figure S1B, gels I, II, and III), implying that RexA also binds dsDNA. Further examination by filter binding shows that RexA preferentially associates with dsDNA compared to single-strand DNA (ssDNA) (Figure 8C, O_R_1-O_R_2 vs. ssO_R_1-O_R_2) but lacks sequence specificity for the λ operator region (O_R_1-O_R_2 vs. O_R_1-O_R_2 scrambled). RexA and CI form ternary complexes on both *O*_L_ and *O*_R_ operator DNA substrates with no significant changes to the overall elution profile of the individual components (Figure 8B), signifying that the abilities of RexA to bind CI and DNA are not mutually exclusive.

We next assessed whether CI oligomerization is a prerequisite for RexA binding. Previous biochemical and crystallographic studies showed that the CI missense mutation D197G (Supporting Information Figure S2A) yields stable CI dimers that retain the ability to bind single operator sites but are impaired in higher order assembly and cooperative repressor functions (Stayrook et al., 2008; Whipple, Kuldell, Cheatham, & Hochschild, 1994). Purified CI D197G dimers form stable complexes with *O*_L_1-*O*_L_2 operator DNA substrates (Supporting Information Figure S2B, gels I, IV, and V). However, we observe no significant shift in either protein when CI D197G is incubated with RexA (Figure S2B, gels II, IV, and VI). When operator DNA substrates are added to this mixture, the elution fractions overlay with positions of the individual RexA-DNA and CI D197G-DNA complexes (Figure S2B, gels I, III, V, and VII), arguing against the formation of a ternary complex. No stable complexes are formed between RexA and either the purified CI NTD or CTD domains, respectively (Figure S2C). Together these data argue that RexA is a non-specific DNA binding protein that can specifically bind assembled CI oligomers.

## Discussion

The two basic components of the phage λ bistable switch are the CI and Cro repressor proteins that bind to operator sites flanking the major lytic promoters, *P*_L_ and *P*_R_, to regulate gene expression. RexA and RexB proteins are certainly not required for the λ bistable switch; however, our results suggest that they are accessory factors able to modulate switch activity, providing an additional layer of regulation that refines the ability of the virus to maintain lysogeny or exit from the lysogenic state. RexA potentiates induction when the inducing signal is low by stabilizing the transition to the lytic state. Some of our experiments show an antagonistic effect of RexB on this RexA-mediated activity (Figures 6,7). Induction is optimal when both RexA and RexB are present (Figure 2), while lysogens mutant for both RexA and RexB are the least prone to transition to the non-immune state (Figures 2, 4, 7). In this way, RexA and RexB act together as two opposing forces to modulate the switch between lysogenic and lytic growth, reducing stochasticity in the switch (Arkin, Ross, & McAdams, 1998; Bednarz, Halliday, Herman, & Golding, 2014; Golding, 2011). Once the bistable switch has transitioned to the non-immune state, our data show that only RexA is necessary to stabilize the lytic configuration and reduce the tendency to return to the immune state (Figure 5). Since RexA is not expressed when the switch is in the nonimmune state, we presume that some *P*_RM_ expression must be required before RexA can act in the experiments shown in Figure 5 (see Figure 9).

**Figure 9.**
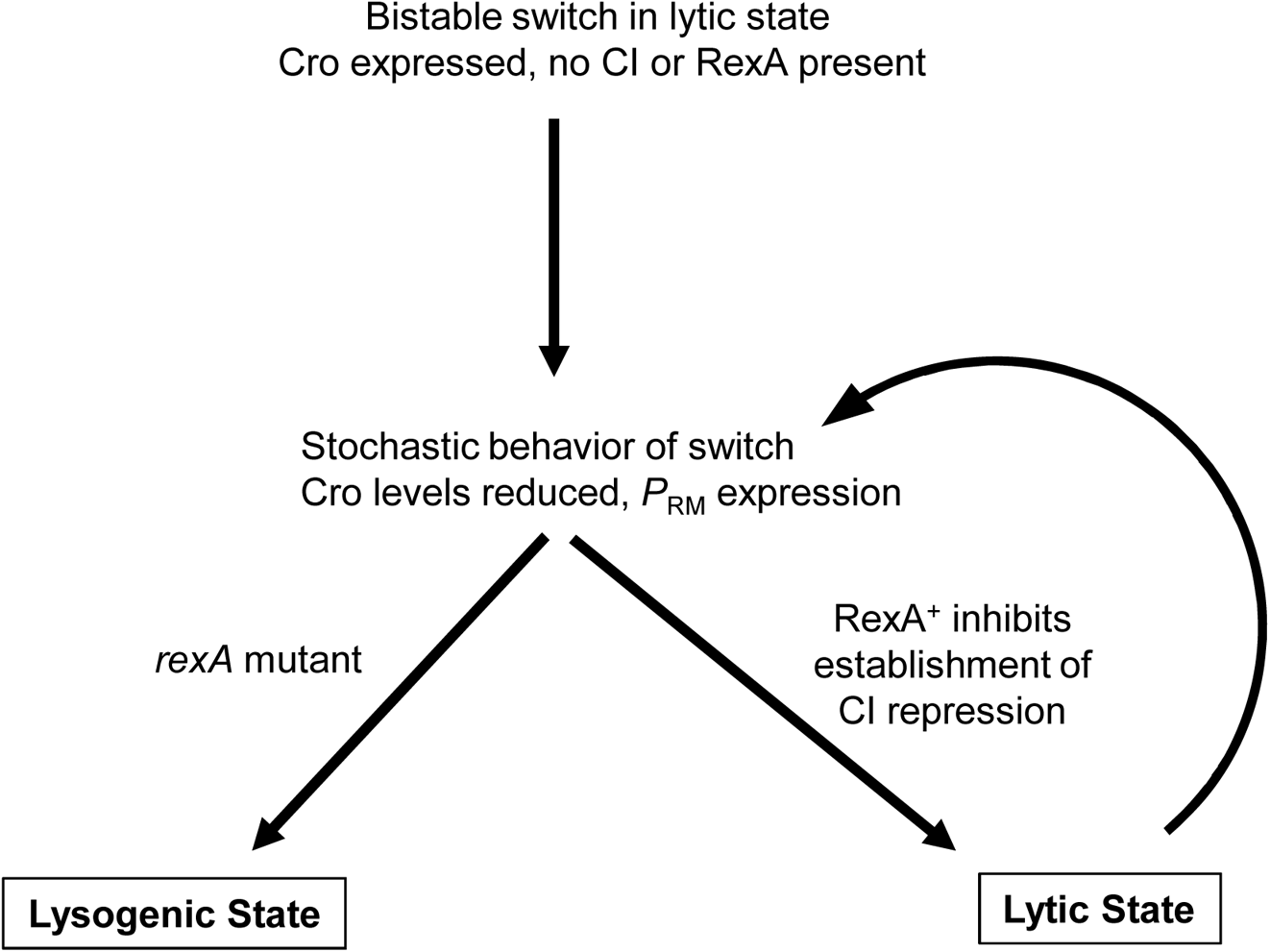
Effect of RexA on the transition from the lytic state to the immune state. When only the phage immunity region is present on the *E. coli* chromosome (see Figure 3), the bistable switch can be in either the immune or nonimmune state. The purified red colonies used for the experiment of Figure 5 have the switch in the nonimmune state, with the Cro protein expressed from *P*_R_ repressing *P*_RM_, so that cI and *rexA* are not expressed. Because of stochastic events, switching to the immune state may occur that relieves Cro repression and allows some *P*_RM_ transcription, resulting in *c*I and *rexA* expression. The data of Figure 5 show that in this situation, RexA protein lessens the probability that CI can establish immune repression, and thus RexA stabilizes the lytic state. If *rexA* is mutant, the switch tends to return to the lysogenic state.

Two types of models may explain Rex effects: Rex proteins either act to directly affect λ behavior, or the Rex proteins alter cell physiology, and this altered physiology in turn affects λ behavior. Our data support the first model, since we demonstrate a direct interaction between the Rex proteins and the phage repressors (Table 2), as well as between the Rex proteins and the phage replication proteins O and Ren (Table S1). Although the Rex system can have energetic effects on the cell, those effects require both Rex proteins (Matz et al., 1982; Parma et al., 1992; Snyder & McWilliams, 1989). Data presented here clearly show that RexA alone exerts an effect on the bistable switch.

Our genetic and biochemical analyses demonstrate that RexA physically interacts with the CI repressor both *in vivo* and *in vitro*. *In vivo*, this interaction was confirmed in the bacterial two-hybrid system as measured by the ability to activate cAMP-dependent promoters. These results suggest that the two proteins can interact via their C-terminal domains. *In vitro*, purified RexA dimers form stable complexes with assembled CI multimers and also bind both dsDNA and ssDNA non-specifically. RexA and CI form ternary complexes on DNA substrates containing λ operator sites, arguing that RexA does not directly inhibit CI intrinsic DNA binding capability, nor does it compete for the same binding sites, which would restrict repressor access. We see no evidence of RexA-mediated CI cleavage analogous to that mediated by RecA protein. We hypothesize that *in vivo*, RexA, facilitated by its interactions with DNA and CI, is present at the λ *P*_L_ and *P*_R_ promotors regulated by CI repressor, where it reduces the ability of CI to establish tight repression, thus pushing the system toward the lytic state (see Figure 10).

**Figure 10.**
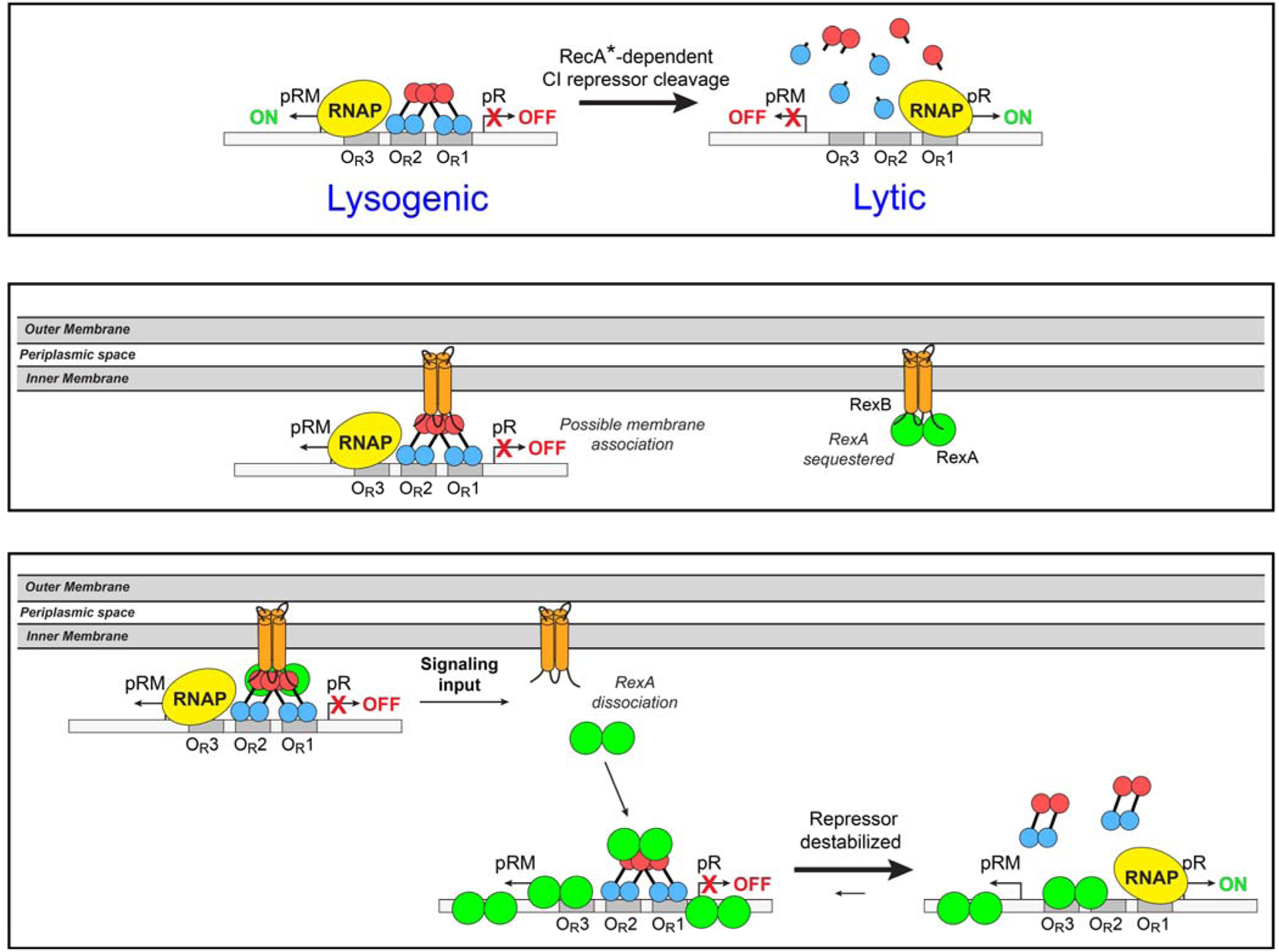
Model for involvement of RexA and RexB in the transition to lytic growth. **Top Panel:** The divergent *P*_RM_ and *P*_R_ promoters and the right operator sites are illustrated. In the lysogenic state, CI dimers are bound cooperatively to *O*_R1_ and *O*_R2_, repressing *P*_R_, and transcription of *P*_RM_ is activated by CI contacting RNA polymerase. Similar CI dimers bound to *O*_L1_ and *O*_L2_ repress the *P*_L_ promoter (not shown in diagram), with long-range DNA looping between the left and right operators mediated by CI repressor molecules. This is the normal state in a λ lysogen. In response to an inducing signal such as DNA damage, RecA protein becomes activated (RecA*) and binds to CI repressor, promoting CI inactivation. This allows transcription from *P*_L_ and *P*_R_ and subsequent Cro expression; Cro will bind to *O*_R3_ and repress *P*_RM_ transcription. **Middle Panel:** Our two-hybrid data suggest protein-protein interactions between CI repressor and the integral membrane protein, RexB. Here, we have shown the CI repressor protein bound to the operator sites while in association with RexB, which may prevent RexA action and deepen repression. This postulated membrane association is not the tight membrane tethering of the lambda genome observed by Hallick and Echols (1973). Our two-hybrid data also show interaction between RexB and RexA (Thomason et al. 2019). **Bottom Panel:** It is possible that CI, RexB, and RexA co-localize at the periphery of the inner membrane to form a complex that includes all three proteins. CI and RexA can be released from RexB, perhaps in response to an environmental signal or a conformational change in RexB. RexA is then free to interact with CI repressor bound to DNA at the operator sites; RexA may also bind DNA nonspecifically. These protein-protein and protein-DNA interactions may destabilize CI repression and activate the lytic state.

RexB is an integral inner membrane protein with four predicted transmembrane segments and three charged cytoplasmic loops; thus, these loops are poised to associate with different binding partners in the cytoplasm (Figure 11) (Krogh, Larsson, von Heijne, & Sonnhammer, 2001; Parma et al., 1992). We find an interaction between RexB and CI repressor in the two-hybrid study (Table 2), suggesting that in a repressed lysogen, CI repressor, bound at its operator sites, may be localized at the inner surface of the cytoplasmic membrane via non-covalent interactions with RexB. CI requires an N-terminal cyclase tag to interact with RexB, however, the cyclase tag on RexB can be at either the N- or C-terminus. Thus, the CI-RexB interaction likely occurs via the RexB central cytoplasmic loop (Figure 11). We previously showed that RexA and RexB physically interact *in vivo* (Thomason et al., 2019); the strength of the RexA-RexB interaction is similar to that of RexA with CI (Table 2). The antagonistic effect of RexB on RexA activity may occur because RexB association with CI bound to the operator sites at the inner membrane makes CI less available to RexA, deepening repression. Alternatively, RexB may titrate RexA, reducing the amount of free RexA protein available to interact with CI and DNA. Regardless of how RexB acts at the molecular level, RexA and RexB must function together to optimize modulation of the CI/Cro bistable switch, with each protein having independent and complementary effects as the phage switches from lysogenic development to lytic growth (see Figure 10).

**Figure 11.**
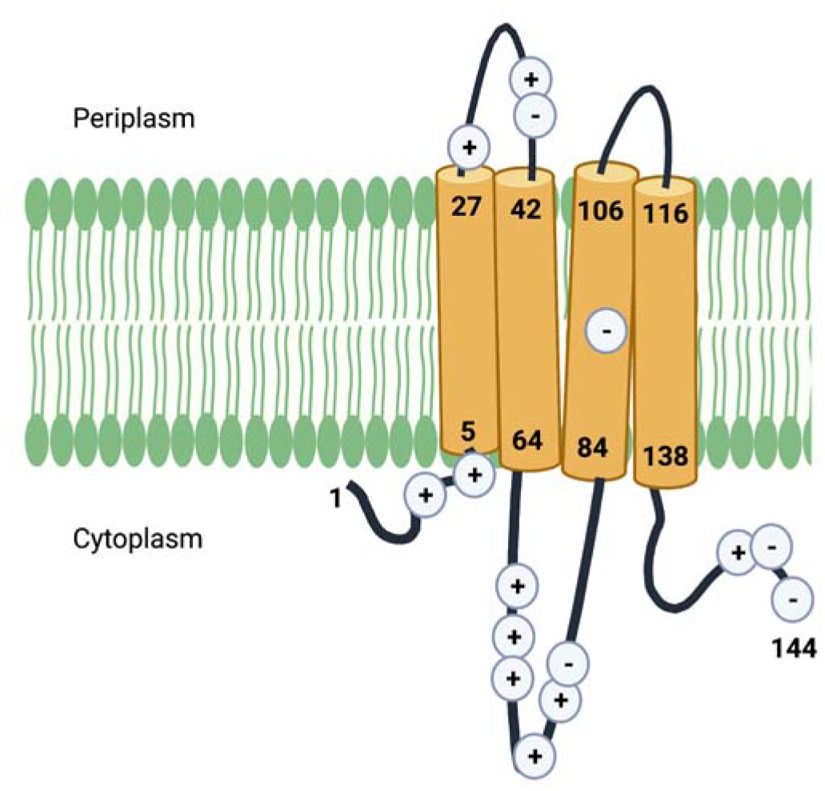
Predicted RexB membrane topology and location of charged amino acid residues. The computer program TMHMM was used to predict the orientation of RexB in the *E. coli* cytoplasmic membrane. The numbers indicate the locations of amino acid residues, with the first and last residues of each transmembrane spanning domain shown. The negatively charged residue in the third transmembrane domain may serve as a proton sink and play some role in energetics. The relative locations of charged amino acids are also indicated. Created with BioRender.com.

RexB continues to be transcribed from *P*_LIT_ after prophage induction as the lytic promoters *P*_L_ and *P*_R_ become active (Fig. 1). The genes encoding the phage DNA replication functions, O and P, are transcribed from *P*_R_. Our two-hybrid analysis (Table S1) suggests a strong physical interaction between the RexB and λ O proteins, consistent with the observation that intracellular RexB stabilizes O protein *in vivo* (MacHattie, 1985; Schoulaker-Schwarz, Dekel-Gorodetsky, & Engelberg-Kulka, 1991). Interaction between RexB and O could tether the phage replication complex to the inner membrane, with RexB serving as a central hub to localize phage DNA replication as well as to coordinate the transition from lysogeny to lytic growth (see Supplementary Results and Discussion).

This work has defined novel and complementary roles for the phage λ RexA and RexB proteins in the switch from lysogeny to lytic growth. Previously, the Rex system was thought to function solely in the lysogenic state to exclude infecting phages such as T4*r*II by triggering cellular energetic defects (Benzer, 1955; Gussin & Peterson, 1972; Matz et al., 1982; Parma et al., 1992). Given its location at the inner membrane, we suspect that the Rex system may sense and respond to the energetic state of the cell, and exclusion may occur as a byproduct.

Despite serving as the basis for pioneering studies in molecular biology and genetics (Benzer, 1955; Crick, Barnett, Brenner, & Watts-Tobin, 1961), the Rex system has long been considered a unique outlier among bacteriophage. The explosion of genome sequencing in recent years, however, has unearthed putative *rex*-like genes in numerous temperate *Mycobacterium* and *Gordonia* phages (Russell & Hatfull, 2017). Homologs from phages Sbash, CarolAnn, and Butters confer a broad viral defense that mirrors the Rex exclusion behavior in λ (Gentile et al., 2019; Mageeney et al., 2020; Montgomery, Guerrero Bustamante, Dedrick, Jacobs-Sera, & Hatfull, 2019). These genes are organized in tandem in a single operon that is often directly adjacent to a CI-like repressor gene, raising the tantalizing possibility that other bacterial viruses also use these analogous systems to fine tune the balance between lysogeny and lytic growth. Future studies will determine the generality of this mechanism and continue the legacy of bacteriophage λ (Golding, 2016) and the Rex system as important experimental platforms.

### Experimental Procedures

#### Materials and Media

Bacterial cultures were grown in L broth containing 10g tryptone, 5g yeast extract and 5g NaCl per liter, and on L plates, which contained ingredients above and 1.5% Difco agar. Cultures for plating phage were grown to exponential phase in tryptone broth containing 10g tryptone, 5g NaCl, and 10mM MgSO_4_ per liter. Phage stocks were maintained in TMG, containing 10mM Tris base, 10 mM MgSO_4_ and 0.01% gelatin, pH7.4. Phage were enumerated on tryptone plates 10g tryptone and 5g NaCl per liter using 0.25 ml of fresh plating cultures mixed with 2.5 ml melted tryptone top agar (0.7% agar) containing 10 g tryptone and 5 g NaCl. MacConkey Lactose agar medium was from Difco and contained 1% lactose and 1.35% agar. Dilutions of bacteria were made in M9 Salts, dilutions of phage were made in TMG.

Bacterial strains used for most experiments are in Table 1, those used for the BACTH analysis are in Table 2. Strains were constructed using a combination of recombineering and P1 transduction methods (Thomason, Costantino, & Court, 2007; Thomason, Sawitzke, Li, Costantino, & Court, 2014). Sequences of single-strand oligonucleotides used for recombineering are available upon request. The mutations to inactivate *rexA* and *rexB* individually or together are described in Thomason et al. (2019). Briefly, most of each gene was removed and replaced with the open reading frame of a selectable drug marker, but two regions were left intact: the distal end of the *rexA* gene, which contains the *P*_LIT_ promoter; and the distal end of the *rexB* gene, which is necessary for optimal transcriptional termination at the intrinsic terminator for the immunity region, *T*_IMM_, which is immediately downstream of *rexB*. For simplicity these mutations are simply indicated in the strain list (Table 1) as *rexA* and *rexB* drug marker replacements. Once the *rex* mutations were inserted in the intact prophage, lysogenic candidates were screened by cross-streaking against a *c*I mutant phage, and those that displayed immunity were further screened with a PCR test (Powell, Rivas, Court, Nakamura, & Turnbough, 1994) to identify monolysogens, which were used for all analyses.

#### UV induction of λ lysogens

Overnight cultures of lysogens and strain A584 for titering released phage were grown in LB at 32 C overnight. Lysogens were diluted 1/500 into 35 ml LB and grown in a 32°C shaking water bath to an OD_600_ of approximately 0.15. Then 30 ml of the cells was harvested by centrifugation and suspended gently in 1 ml of TMG. 29 ml TMG was added, and the wash and suspension steps repeated. After the second wash, 14 ml TMG was added to the cells, for a total of 15 ml, resulting in a two-fold concentration and ~2×10^8^ cells/ml in TMG. The washed lysogenic cells were placed in a sterile empty petri dish and working under a red light, were irradiated with UV for the indicated amount of time. Every 5 seconds, 100 μl was withdrawn and dispensed into a sterile Eppendorf tube, out to 30 sec. 10 microliters from each Eppendorf was added to 10ml LB in a 50ml baffled flask. Cells were incubated in the dark in a 37°C shaking water bath for two hours. Serial dilutions of the resulting lysates were made and 10 μl of each dilution was spotted on a lawn of A584 poured onto TB plates; the dilutions were stored overnight. The next day the approximate titers of each dilution were estimated from the spot plates, and more accurate titers determined as follows: 50 or 100μl of the appropriate phage dilution was mixed with 0.25 ml A584 in a small plating tube, this cell-phage mix remained at room temperature for at least 10 min, then 2.5 ml melted TB top agar was added, and the contents immediately poured on a TB plate. Once the top agar hardened plates were inverted and incubate overnight at 37°C. The next day, the number of plaques on each plate was counted and titers were determined for each UV time point. Plating bacteria were made by diluting the A584 overnight 33-fold into five ml tryptone broth containing 10mM MgSO4, growing for 2.5 hrs at 32°C with aeration, after which five ml of TMG (10mM Tris 10mM MgSO4 10mM gelatin) was added. This thin culture was used to assay released phage.

#### Papillation assays

The papillation of strains carrying the dual *P*_L_ *P*_R_ reporters with a *c*I*857 ind1* allele and *rexA* and *rexB* mutations was examined by plating appropriate dilutions of fresh LB overnight cultures on MacConkey-Lactose to obtain isolated single colonies. MacConkey-Lactose petri plates were incubated at 32-34°C until papillae arose within individual colonies (Fig. 4). All plates in a set were incubated under identical conditions. Once papillae developed, the plates were photographed, and the pointed end of a small pipette tip was used to pick individual papillae and purify them to single red colonies on MacConkey-Lactose; these plates were incubated at 30°C degrees. Twenty-four independent red colony isolates of each genotype were propagated in this fashion for the RecA^+^ strains, and eighteen for the RecA^−^. Once these red colonies were purified away from white colonies, overnight cultures from the red colonies were grown in L broth at 30°C and plated on MacConkey-Lactose solid agar for single colonies. After overnight incubation at 30°C, colonies were counted for each independent culture and the number of red and white colonies tabulated. The percentage of white colonies among reds was determined for each genotype (Fig. 5). In every case, a few colonies of each genotype failed to give any white colonies. Such non-reverting red colonies likely contain mutations in the *c*I repressor gene, which generate inactive CI repressor protein; these were not included in the analysis. The final number of colonies analyzed is indicated in the legend to Fig. 5. Numerical data are shown in Supporting Information Table S4; these data were used to generate the bar graphs shown in Fig. 5A and 5B.

#### Kinetics of induction monitoring P_L_ N-luc reporter

Overnight cultures of strains LT1657, LT1659, LT1895, and LT1897 were grown in L broth from single colonies. Cultures were diluted 500-fold into 25ml L broth in 125 ml baffled flasks in a 37°C shaking water-bath. After 1.5 hrs, 1.0 ml samples were withdrawn from the flasks and 3ng/ml Mitomycin C (MC) was added to each flask. For these and subsequent samples, taken every 30 min, the A600 was determined, and luciferase assays were performed immediately, using the Promega Luciferase Assay System (catalog no. E1500) according to the company’s directions. Culture aliquots in L broth (0.1ml) were added directly to 0.4 ml of Cell Culture Lysis Reagent with 2.5mg/ml BSA and 1.25 mg/ml lysozyme. Cell lysate (25μl) was mixed with 0.1ml of luciferase substrate (Promega Luciferase Assay Reagent, catalog no. E151A), incubated for 2.0 min and read in a BD Pharmingen Monolight 3010 single sample luminometer for 10s. Each sample was assayed in duplicate. The relative light unit (RLU) was normalized to the A600. Three biological replicates of the experiment were performed, one representative experiment is shown (Figure 6).

#### Analysis of spontaneous phage release in lysogenic cultures

Lysogenic cells were first washed to remove viral particles: five ml overnight cultures of lysogenic strains were pelleted in the Sorvall at 6700rpm for 7min. Pellets were suspended in 1ml TMG, additional TMG was added to ~25ml total volume, and suspensions were mixed by vortexing. Cells were again pelleted, suspended in TMG, and pelleted as before, then the cell pellet was suspended at the original 5ml volume and diluted 1/400 into 10ml L broth in 125ml baffled flasks. Aliquots (1ml) of each genotype were removed and filtered with a 0.2μm filter; 0.3ml of the filtered lysate was plated on tryptone plates using C600 as host to determine initial phage titer, which was negligible in all cases. Cultures were incubated with shaking in a 32°C water bath until the OD_600_ was 0.2-0.3. Final OD_600_ was determined for each culture and the number of viable cells per ml was determined from colony counts obtained by plating dilutions of lysogenic cultures on L agar petri plates and incubating plates overnight at 32°C. 1.5ml each culture was removed and filtered with a 0.2μm filter, three 10-fold serial dilutions into TMG were made for each of the filtered lysates. 0.25ml of C600 plating culture was mixed with 2.5ml melted tryptone top agar and poured onto tryptone petri plates. Once plates hardened, 10μl of the undiluted lysate and each dilution was spotted on the surface of the lawn. Plaques in each 10μl spot were counted to determine approximate titers of the cultures. These approximations were used to guide the selection of appropriate dilutions of filtered lysates to determine an accurate number of phage particles in the supernatants. Both clear (*c^−^*) and turbid (*c^+^*) plaques were enumerated. Clear plaques result from mutations in the phage that prevent lysogeny and accounted for less than 5% of the total phage yield for all genotypes. Numerical data are shown in Supporting Information Table S5.

#### Two-hybrid analysis

The Bacterial Adenylate Cyclase Two Hybrid System Kit (BACTH Kit) from Euromedex (www.euromedex.com) was used to look for protein-protein interactions between the RexA and RexB proteins and the two phage repressor proteins in *E. coli*. Construction of *rexA* and *rexB* plasmids was previously described (Thomason 2019). Recombineering was used to insert the *c*I and *cro* repressor genes into the four two-hybrid vectors: the low-copy KanR pKT25 and pKNT25 plasmids with the T25 adenylate cyclase domain on the N- and C-terminal domains fused in frame to the gene of interest, respectively. The two repressor genes were also inserted into the high-copy AmpR pUT18 and pUT18C plasmids, with the T18 adenylate cyclase domain fused in frame to the C- and N-terminal domains of the gene of interest, respectively. For BACTH complementation assays, various pairs of T25 (KanR) and T18 (AmpR) plasmids were introduced into the *cya* mutant strain BTH101 by co-electroporation. The BTH101 derivatives containing these plasmid pairs are listed in Table S2 A-E of the Supporting Information. After outgrowth, 10μl drops of ten-fold serial dilutions of the transformed cells were spotted onto petri plates containing L agar with kanamycin (30μg/ml), ampicillin (100μg/ml) and X-gal (50μg/ml). Plates were incubated at 32°C overnight. Colonies from these petri plates were subsequently purified on M63 minimal maltose solid agar containing the same drugs plus 1mM IPTG. Growth on minimal maltose indicated cAMP-dependent sugar utilization and thus a positive interaction between the two phage proteins being tested; such interaction results in interaction of the two fused adenylate cyclase domains (Supporting Information Table S1). Out of 51 strains tested, 13 grew on minimal maltose, and β-galactosidase assays were used to provide an estimate of the strength of interaction for these pairs of plasmid-borne phage proteins (Table 2). For these assays, colonies picked from the minimal maltose agar plates were used to generate overnight cultures grown in L broth containing the same antibiotics at the same concentrations. The day of the assay, cells were diluted 100-fold into L broth lacking antibiotics and β-galactosidase assays were done according to Thomason et al. (2019). At least three independent cultures were measured for each plasmid pair. A similar protocol was followed for the phage O, P, and Ren proteins (see Supporting Information Results and Discussion and Table S1).

#### Cloning, expression, and purification of bacteriophage λ RexA and CI constructs

DNA encoding the full-length bacteriophage λ RexA (UniProt P68924) and CI (UniProt P03034) proteins were codon optimized for *E. coli* expression and synthesized commercially by Bio Basic Inc. The DNA encoding full-length RexA (residues 1-279) was amplified by PCR and cloned into pET21b, introducing a 6xHis tag at the C-terminus. DNA encoding full-length CI (residues 1-237), as well as its N-terminal DNA binding domain (NTD, residues 1-93) and the C-terminal multimerization domain (CTD, residues 133-237), was separately amplified by PCR and cloned into pET21b. The D197G mutation was introduced into full-length CI via the Quikchange Mutagenesis Kit (Agilent). All RexA and CI constructs were transformed into BL21(DE3) cells, grown at 37°C in Terrific Broth to an OD600 of 0.7-0.9, and then induced with 0.3 mM IPTG overnight at 19°C. Cells were pelleted, washed with nickel loading buffer (20 mM HEPES pH 7.5, 500 mM NaCl, 30 mM imidazole, 5% glycerol (v:v), and 5 mM β-mercaptoethanol), and pelleted a second time. Pellets were typically frozen in liquid nitrogen and stored at −80°C for later use.

Thawed 500 ml pellets of each RexA and CI construct were resuspended in 30 ml of nickel loading buffer supplemented with 5 mg DNase, 5 mM MgCl_2_, 10 mM PMSF, and a Roche complete protease inhibitor cocktail tablet. Lysozyme was then added to a concentration of 1 mg/ml and the mixture was incubated for 10 minutes rocking at 4°C. Cells were disrupted by sonication and the lysate was cleared via centrifugation at 13 000 rpm (19 685 g) for 30 minutes at 4°C. The supernatants were each filtered through a 0.45 μm filter, loaded onto a 5 ml HiTrap chelating column charged with NiSO_4_, washed with nickel loading buffer, and then eluted via an imidazole gradient from 30 mM to 1 M. Pooled RexA fractions were further dialyzed overnight at 4°C into S loading buffer (20 mM HEPES pH 7.5, 50 mM NaCl, 1 mM EDTA, 5% glycerol (v:v), and 1 mM DTT) and then loaded onto a 5 ml HiTrap SP column equilibrated with S loading buffer. The SP column was washed in the same buffer and then RexA was eluted with a NaCl gradient from 50 mM to 1 M. Peak fractions from the SP column were pooled, concentrated, and further purified by size exclusion chromatography (SEC) using a Superdex 75 16/600 pg column. RexA protein was exchanged into a final buffer of 20mM HEPES pH 7.5, 150mM KCl, 5 mM MgCl_2_, and 1mM DTT during SEC and concentrated to 10-70 mg/ml.

The chelating column elution fractions containing CI protein were pooled, concentrated, and further purified directly by SEC using either a Superdex 200 16/600 pg column (full-length wildtype) or Superdex 75 16/600 pg column (D197G mutant, NTD, and CTD). CI proteins were exchanged into a final buffer of 20mM HEPES pH 7.5, 150mM KCl, 5 mM MgCl_2_, and 1mM DTT during SEC and concentrated to 10-40 mg/ml.

#### Size exclusion chromatography coupled to multi-angle light scattering (SEC-MALS)

Purified RexA at 4 mg/mL was subjected to size-exclusion chromatography (SEC) using a Superdex 75 10/300 column (GE) equilibrated in SEC buffer (20 mM HEPES pH 7.5, 150 mM KCl, 5 mM MgCl_2_, and 1mM DTT). The column was coupled to a static 18-angle light scattering detector (DAWN HELEOS-II) and a refractive index detector (Optilab T-rEX) (Wyatt Technology). Data were collected continuously at a flow rate of 0.5 mL/min. Data analysis was carried out using the program Astra VI. Monomeric BSA at 4 mg/mL (Sigma) was used for normalization of the light scattering detectors and data quality control.

#### Preparation of DNA substrates from oligonucleotides

All DNA oligonucleotides were synthesized commercially by Integrated DNA Technologies (IDT). Lyophilized single-stranded oligonucleotides were resuspended to 1 mM in 10 mM Tris-HCl and 1 mM EDTA and stored at minus 20°C until needed. For filter binding, single-stranded oligonucleotides were 5’ end-labeled with [γ32P]ATP using polynucleotide kinase (New England Biolabs) and then purified on a P-30 spin column (BioRad) to remove unincorporated label. Duplex DNA substrates were prepared by heating equimolar concentrations of complementary strands (denoted as ‘us’ and ‘ls’ indicating upper and lower strands) to 95°C for 15 minutes followed by cooling to room temperature overnight and then purification on an S-300 spin column (GE) to remove single stranded DNA. Sequences for all substrates can be found in Supporting Information Table S3.

#### Analytical size exclusion chromatography (SEC)

50 μl samples containing full-length CI protein at 125 μM, RexA protein at 125 μM, or a 1:1 mixture of both proteins was incubated for 15 minutes at room temperature and analyzed by SEC using a Superdex 200 PC 3.2 column (GE Healthcare) equilibrated in SEC buffer (20 mM HEPES, pH7.5, 150 mM KCl, 5 mM MgCl_2_, and 1mM DTT). To assess CI and RexA interactions with DNA, samples containing the annealed double-stranded DNA substrates OR_1_-OR_2_ or OL_1_-OL_2_ were also prepared at 2:1.2 protein to DNA molar ratio. Individual DNA substrates were injected alone at a concentration of 75 μM for comparison. All eluted fractions were further analyzed by SDS-PAGE using 4 –20% gradient gels, and then silver-stained to visualize DNA and Coomassie-stained to visualize protein. Samples were similarly prepared for the D197G, NTD, and CTD CI constructs but were analyzed using a Superdex 75 PC 3.2 column (GE Healthcare) equilibrated in SEC buffer.

#### Filter binding

Filter binding assays were carried out in buffer containing 25 mM MES (pH 6.5), 2.0 mM MgCl2, 0.1 mM DTT, 0.01 mM EDTA, and 40 μg/mL BSA. Binding was performed with wildtype RexA at 30°C for 10 min in a 30 μL reaction mixture containing 14.5 nM unlabeled DNA and 0.5 nM labelled DNA. Samples were filtered through KOH-treated nitrocellulose filters (Whatman Protran BA 85, 0.45 μm) using a Hoefer FH225V filtration device for approximately 1 min. Filters were subsequently analyzed by scintillation counting on a 2910TR digital, liquid scintillation counter (PerkinElmer). All measured values represent the average of at least three independent experiments (mean ± standard deviation) and were compared to a negative control to determine fraction bound.

## Acknowledgements

We thank Brenda Shafer for expert technical assistance, Alison Rattray for statistical analyses, and Nina Costantino for critical reading of the manuscript. L.C.T. thanks Nathan Brown for insightful discussions about the Rex system. We also thank Richard Fredrickson and Jonathan Summers at NCI-Frederick Scientific Publications, Graphics, and Media for expert assistance with photography, and M. Spencer, N. Shrader, T. Hartley, and K. Pike from the CRTP Genomics Laboratory of the Frederick National Lab for Sanger sequencing. This work was supported, in part, by the Intramural Research Program of the National Institutes of Health, National Cancer Institute, Center for Cancer Research. This project has been funded in whole or in part with federal funds from the National Cancer Institute, National Institutes of Health, under contract HHSN26120080001E. The content of this publication does not necessarily reflect the views or policies of the Department of Health and Human Services, nor does mention of trade names, commercial products, or organizations imply endorsement by the U.S. Government. This research was supported [in part] by the Intramural Research Program of the NIH, National Cancer Institute, Center for Cancer Research. M.C.A. is supported by a NIFA predoctoral fellowship (2020-67034-31750).

## Author Contributions

L.C.T. performed genetic experiments and wrote the manuscript, C.C. performed genetic experiments. C.J.S., C.J.H., and M.C.A. performed biochemistry experiments, J.S.C. designed biochemical experiments, provided guidance and financial support, and edited the manuscript. D.L.C. provided financial support and guidance and edited the manuscript.

## Data Availability Statement

The data that support the findings of this study are available from the corresponding author upon reasonable request.

