## Supplemental for "Bacteriophage λ RexA and RexB Functions Assist the Transition from Lysogeny to Lytic Growth"

**Inventory of Supporting Information:**

**Supporting Results and Discussion: The RexA and RexB proteins interact with phage λ replication protein O and Ren protein.**

**Table S1. β-galactosidase measurements for BACTH system RexA/RexB with λ O and Ren**

**Table S2. Bacterial Strain Growth on Minimal Maltose for Bacterial Adenylate Cyclase Two-Hybrid (BACTH) with RexA, RexB, CI and Cro plasmids**

**A. RexA by CI**

**B. RexA by Cro**

**C. RexB by CI**

**D. RexB by Cro**

**E. CI by Cro**

**Table S3. DNA oligonucleotide substrates**

**Table S4. Data for Fig. 5: Propensity of the bistable switch to return to the immune state from the nonimmune state**

**Table S5. Data for Fig. 6: Dependence of phage release on *rexAB* genotype in lysogenic strains growing in liquid culture**

**Figure S1. Disk diffusion assays of RecA+ Cro+ *c*I*857* *ind1* strains challenged with Mitomycin C on MacConkey Lactose indicator media.**

**Figure S2. SEC analysis of CI D197G mutant and truncation constructs**

**Supporting References**

**Supporting Results and Discussion**

**The RexA and RexB proteins interact with phage λ replication proteins O and Ren**. Phage λ encodes two genes known to be involved in phage DNA replication, *O* and *P*; together these genes are classified as an “Initiator-helicase loader” replication module (Weigel & Seitz, 2006). The λ *O* gene encodes the replication initiation protein, O, an analog of *E. coli* DnaA. O protein binds to the phage origin of DNA replication, embedded within the *O* gene. The downstream gene, *P*, encodes the P protein, which recruits the *E. coli* replication complex to the phage origin via interaction with host DnaB helicase (Chase et al., 2018). A third gene, *ren*, is located beyond *P* such that the three genes *O*, *P*, and *ren* are terminally overlapping and co-translated. All phages with “Initiator-helicase loader” type replication modules also have a *ren* gene (Weigel & Seitz, 2006), implying some as yet unidentified role for Ren in phage replication.

Results from several groups suggest a connection between the Rex system, phage λ DNA replication, and *ren* function. Toothman and Herskowitz (1980) showed that heteroimmune λ phage, which are not subject to CI repression, grow poorly in Rex^+^ lysogens when mutant for *ren*; they identified genetic suppressors of this inhibition in the *O* and *P* genes. MacHattie (1985) found that RexB expression improves the establishment of plasmids undergoing λ OP-dependent DNA replication, while Schoulaker-Schwarz, Dekel-Gorodetsky, and Engelberg-Kulka (1991) found that RexB protein expression *in vivo* increases the stability of λ O protein. Additionally, a yeast two-hybrid analysis (Rajagopala, Casjens, & Uetz, 2011) showed a direct interaction between RexB and Ren proteins.

In light of these clues that Rex function may also impact phage replication once the transition to lytic growth has been achieved, and since RexB continues to be expressed during lytic growth from the *P*_LIT_ promoter (Thomason et al., 2019), we used the bacterial two hybrid system (BATCH) to look for possible interactions between the Rex proteins and O, P, and Ren. We find that the affinity between RexB and O is very high (Table S1) and that both RexB and RexA interact with O and Ren (Table S1). In contrast, when RexB and RexA were tested against the λ P protein, no growth was found on minimal maltose, arguing that there is no physical interaction between RexB or RexA and λ P protein.

While an understanding of the effects on the Rex system on phage replication and elucidation of Ren protein function is beyond the scope of this paper, our results, in combination with those of others, do suggest that there is more to learn here. We suspect that the Rex system may act as a pivot point between λ lysogenic development and lytic growth, and that the membrane protein RexB may enhance establishment of DNA replication for the phage by interacting with the replication initiator protein O at the inner surface of the cytoplasmic membrane (see Figure S1). This would provide a specific molecular mechanism for the observed membrane tethering of the phage DNA observed by Hallick and Echols (1973). The mechanism of switching from the early circle-to-circle mode of DNA replication to the rolling circle mode is still not understood; it is possible that the Rex system and Ren may be involved in this transition.

**Table S1. β-galactosidase measurements for BACTH system RexA/RexB with λ O and Ren^†^**

| **Strain Number** | **Relevant Genotype** | **Configurations of hybrid proteins** | **β-galactosidase Units^‡^** |
| --- | --- | --- | --- |
| LT2212 | BTH101[pKT25-*rexA*] [pUT18C-*O*] | *cya25-rexA*  *cya18-O* | 458 ± 10 |
| LT2213 | BTH101[pKNT25-*rexA*] [pUT18C-*O*] | *rexA-cya25*  *cya18-O* | 299 ± 126 |
| LT2201 | BTH101[pKNT25-*rexA*] [pUT18-*O*] | *rexA-cya25*  *O-cya18* | 125 ± 37 |
| LT2206 | BTH101 [pKT25-*rexB*] [pUT18C-*O*] | *cya25-rexB*  *cya18-O* | 1883 ± 79 |
| LT2205 | BTH101[pKNT25-*rexB*] [pUT18-*O*] | *rexB-cya25*  *O-cya18* | 1099 ± 226 |
| LT2204 | BTH101[pKT25-*rexB*] [pUT18-*O*] | *cya25-rexB*  *O-cya18* | 492 ± 20 |
| LT2211 | BTH101[pKNT25-*rexA*] [pUT18C-*ren*] | *rexA-cya25*  *cya18-ren* | 329 ± 61 |
| LT2203 | BTH101 [pKNT25-*rexB*] [pUT18-*ren*] | *rexB-cya25*  *ren-cya18* | 188 ± 4 |
| LT2208 | BTH101[pKT25-*rexB*] [pUT18C-*ren*] | *cya25-rexB*  *cya18-ren* | 107 ± 44 |

^†^Only plasmid pairs that gave β-gal measurements over 100 units are shown.

^‡^In all cases, the experiment was repeated three times with two technical replicates. Standard error of the mean (s.e.m.) is shown.

**Table S2. Bacterial Strain Growth on Minimal Maltose for Bacterial Adenylate Cyclase Two-Hybrid (BACTH) with RexA, RexB, CI and Cro plasmids**

**A. RexA by CI**

| **Strain** | **Relevant Genotype^†^** | **Configurations of hybrid proteins** | **Growth on Minimal Maltose**  **X-gal** |
| --- | --- | --- | --- |
| LT2333 | BTH101[pKT25-*rexA*] [pUT18C-*c*I] | *cya25-rexA*  *cya18*-*c*I | Yes |
| LT2334 | BTH101[pKNT25-*rexA*] [pUT18C-*c*I] | *rexA-cya25*  *cya18-c*I | No |
| LT2379 | BTH101[pKT25-*c*I] [pUT18-*rexA*] | *cya25*-*c*I  *rexA-cya18* | No |
| LT2380 | BTH101[pKNT25-*c*I] [pUT18-*rexA*] | *c*I- *cya25*  *rexA-cya18* | No |
| LT2381 | BTH101[pKT25-*c*I] [pUT18C-*rexA*] | *cya25-c*I  *cya18*-*rexA* | Yes |
| LT2382 | BTH101[pKNT25-*c*I] [pUT18C-*rexA*] | *cI-cya25*  *rexA-cya18* | No |
| LT2391 | BTH101[pKT25-*rexA*] [pUT18-*c*I] | *cya25-rexA*  *cI-cya18* | No |
| LT2392 | BTH101[pKNT25-*rexA*] [pUT18-*c*I] | *rexA-cya25*  *c*I*-cya18* | No |
| LT2447 | LT2333 *nadA::Tn*10 (λ*c*I<>*kan Δ{N-int}*) | *cya25-rexA*  *cya18*-*c*I | Yes |
| LT2448 | LT2381 *nadA::Tn*10 (λ*c*I<>*kan Δ{N-int}*) | *cya25-c*I  *cya18*-*rexA* | No |
| LT2450 | LT2391 *nadA::Tn*10 (λ*c*I<>*kan Δ{N-int}*) | *cya25-rexA*  *cI-cya18* | No |

4

3

2

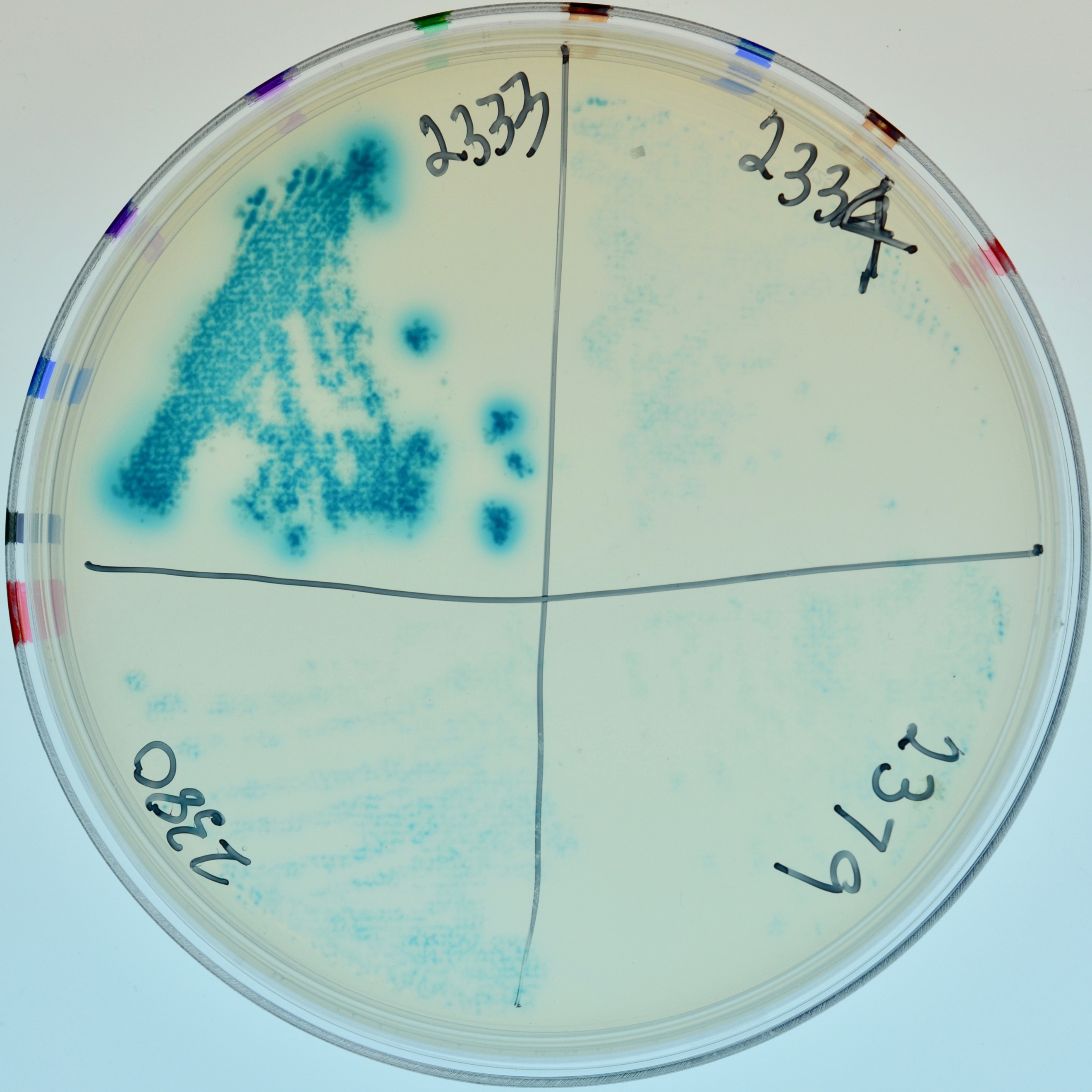

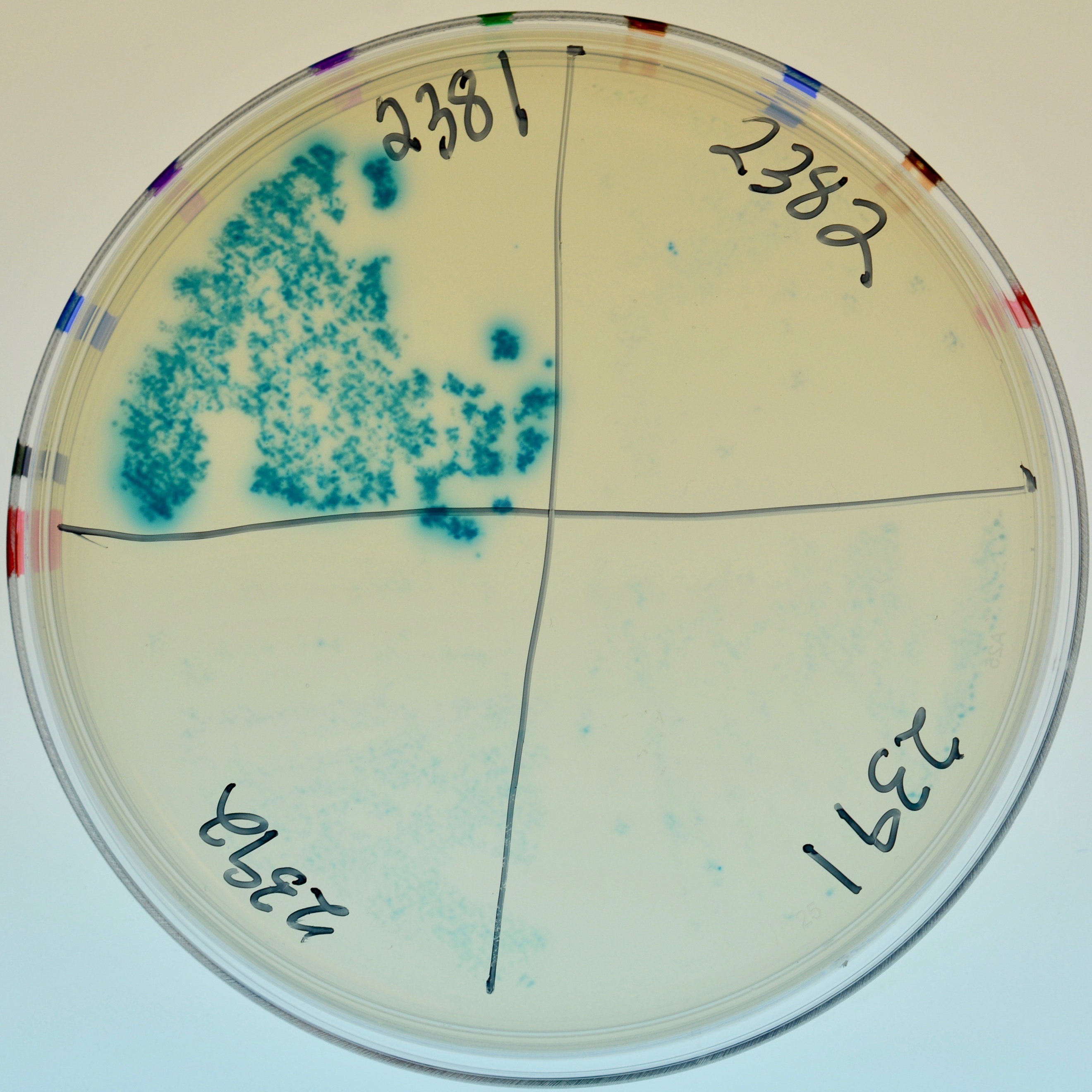

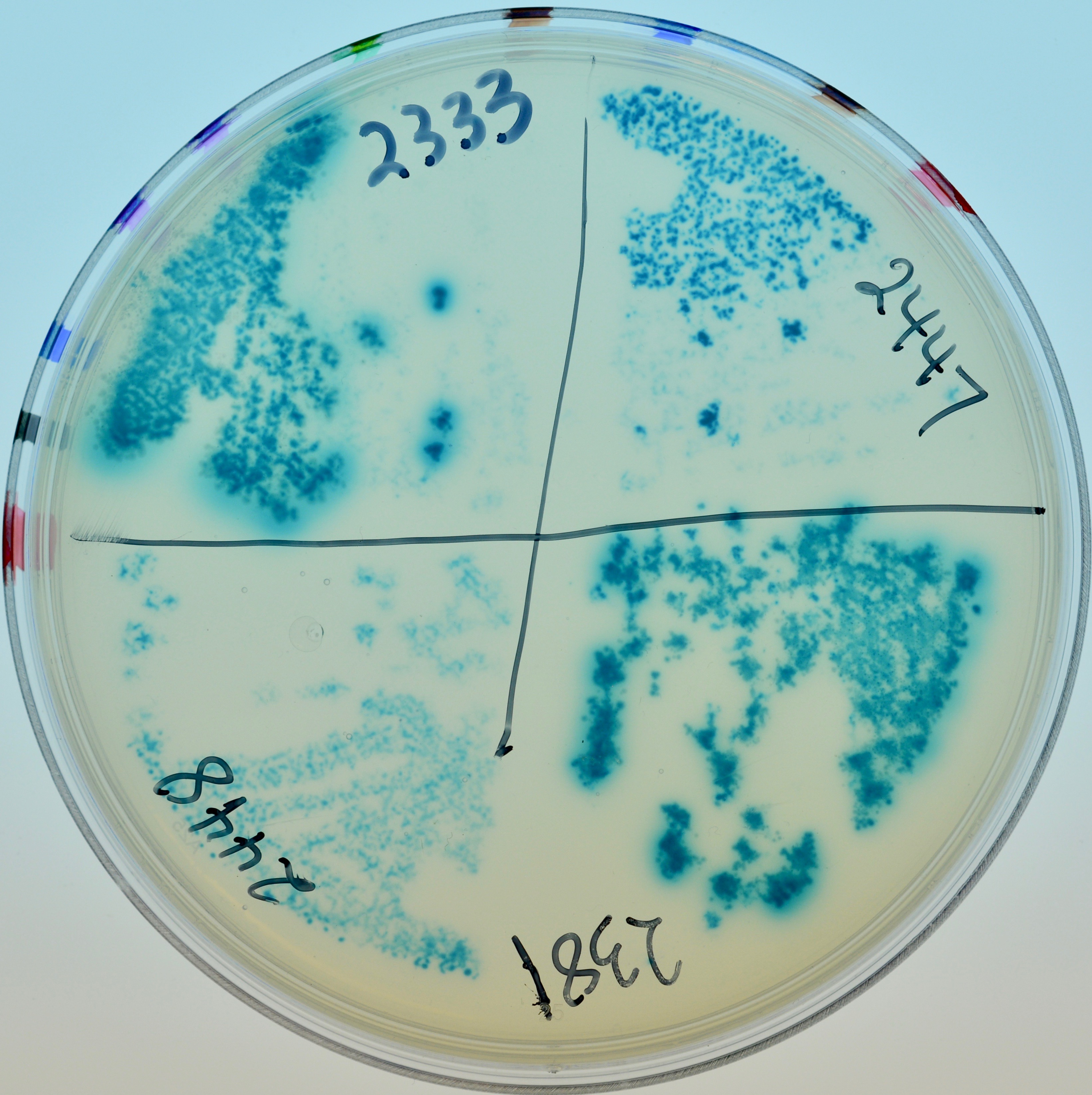

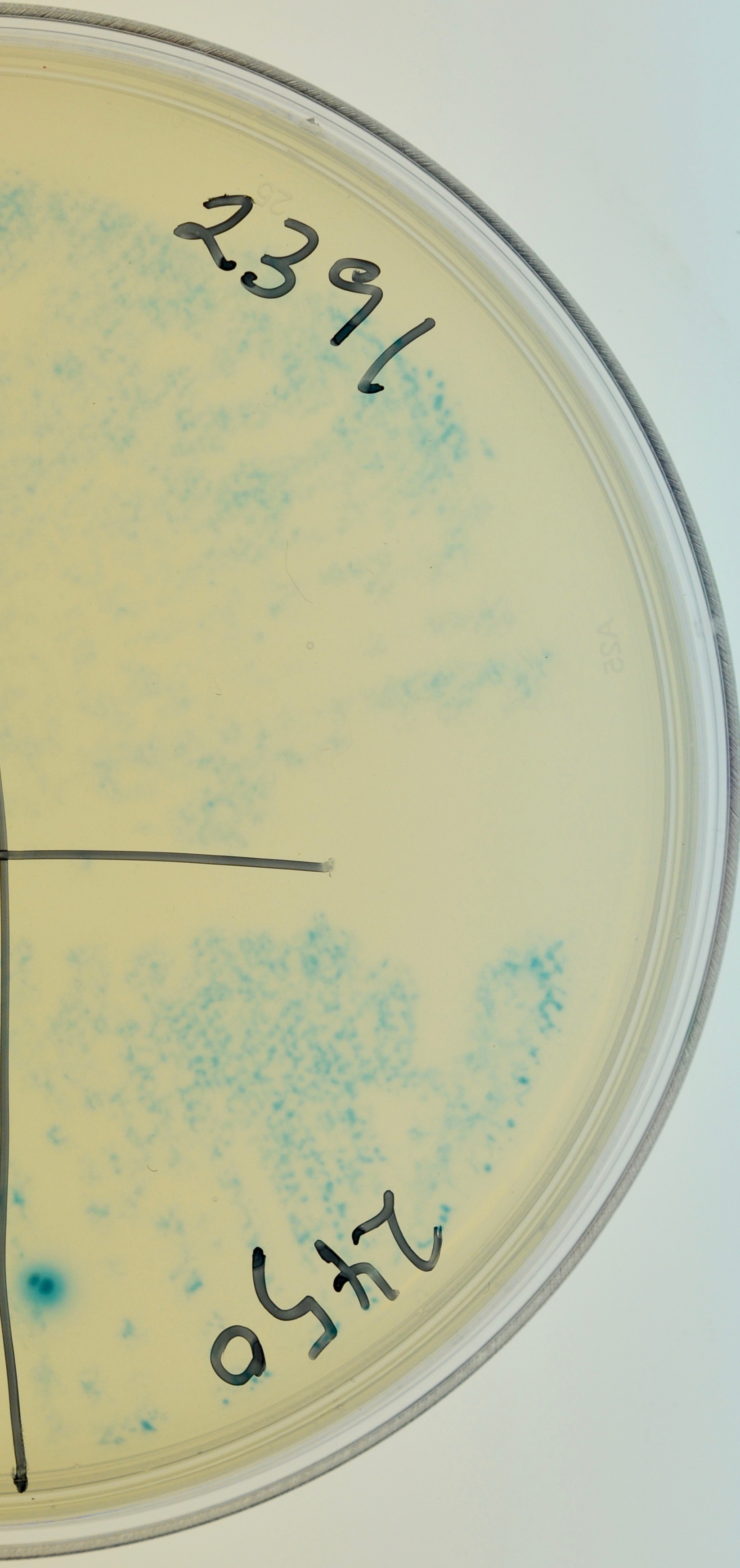

1

Appearance of colonies on minimal maltose with Amp100, Kan30, IPTG and X-gal.

1. From top left, clockwise: LT2333, LT2334, LT2379, LT2380.

2. From top left, clockwise: LT2381, LT2382, LT2391, LT2392

3. From top left, clockwise: LT2333, LT2447, LT2381, LT2448.

4. Top, LT2391; bottom, LT2450.

**B. RexA by Cro**

| **Strain** | **Relevant Genotype^†^** | **Configurations of hybrid proteins** | **Growth on Minimal Maltose**  **X-gal** |
| --- | --- | --- | --- |
| LT2387 | BTH101[pKT25-*cro*] [pUT18-*rexA*] | *cya25*-*cro*  *rexA-cya18* | No |
| LT2388 | BTH101[pKT25-*cro*] [pUT18C-*rexA*] | *cya25-cro*  *cya18*-*rexA* | Yes |
| LT2389 | BTH101[pKT25-*rexA*] [pUT18-*cro*] | *cya25-rexA*  *cro-cya18* | No |
| LT2390 | BTH101[pKNT25-*rexA*] [pUT18-*cro*] | *rexA-cya25*  *cro-cya18* | No |
| LT2399 | BTH101[pKT25-*rexA*] [pUT18C-*cro*] | *cya25-rexA*  *cya18-cro* | No |
| LT2400 | BTH101[pKNT25-*rexA*] [pUT18C-*cro*] | *rexA-cya25*  *cya18-cro* | No |
| LT2401 | BTH101[pKNT25-*cro*] [pUT18-*rexA*] | *cro-cya25*  *rexA-cya18* | No |
| LT2402 | BTH101[pKNT25-*cro*] [pUT18C-*rexA*] | *cro-cya25*  *cya18-rexA* | No |
| LT2449 | LT2388 *nadA::Tn*10 (λ*c*I<>*kan Δ{N-int})* | *cya25-cro*  *cya18-rexA* | No |
| LT2451 | LT2399 *nadA::Tn*10 (λ*c*I<>*kan Δ{N-int}*) | *cya25-rexA*  *cya18-cro* | No |

3

2

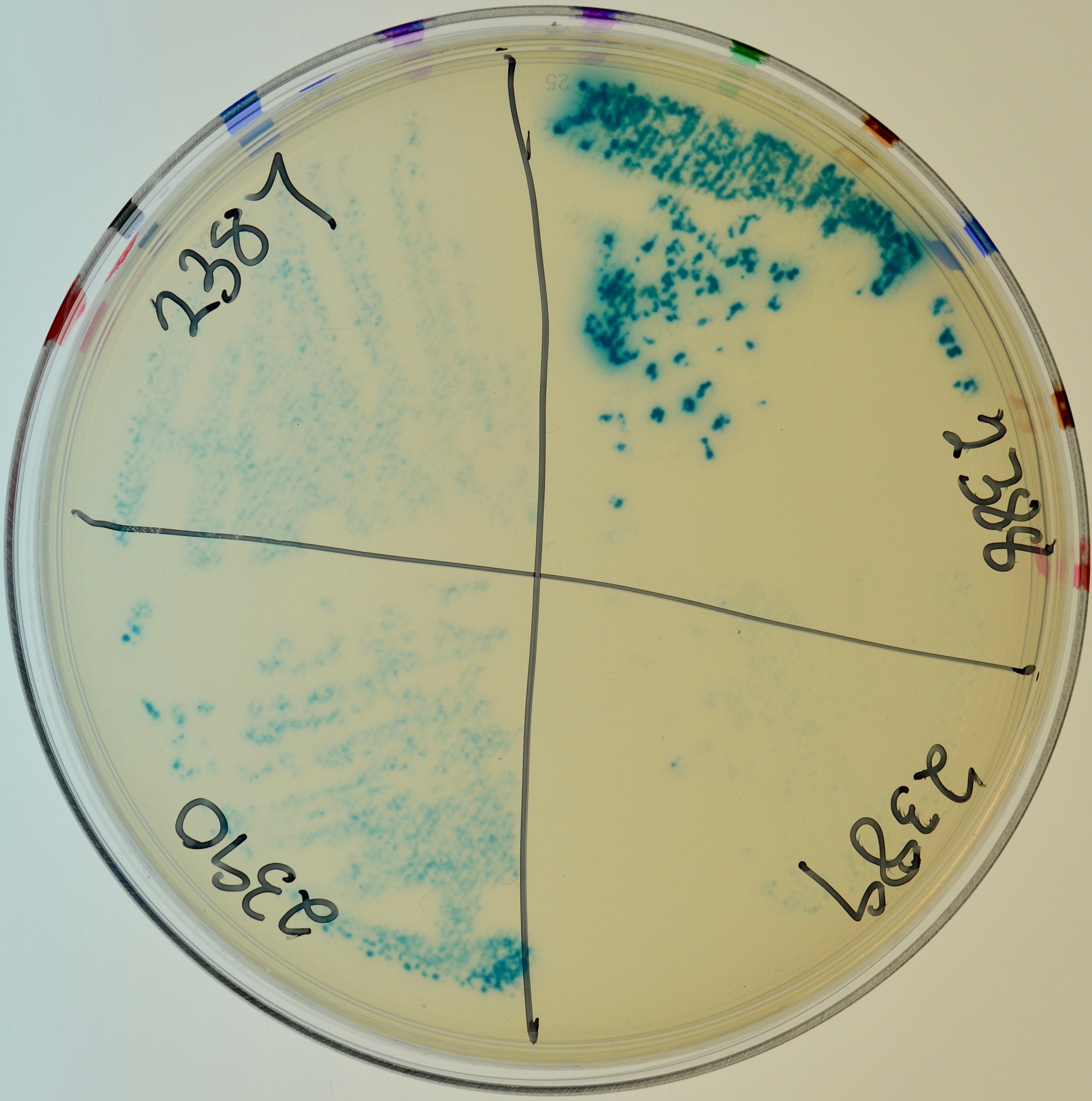

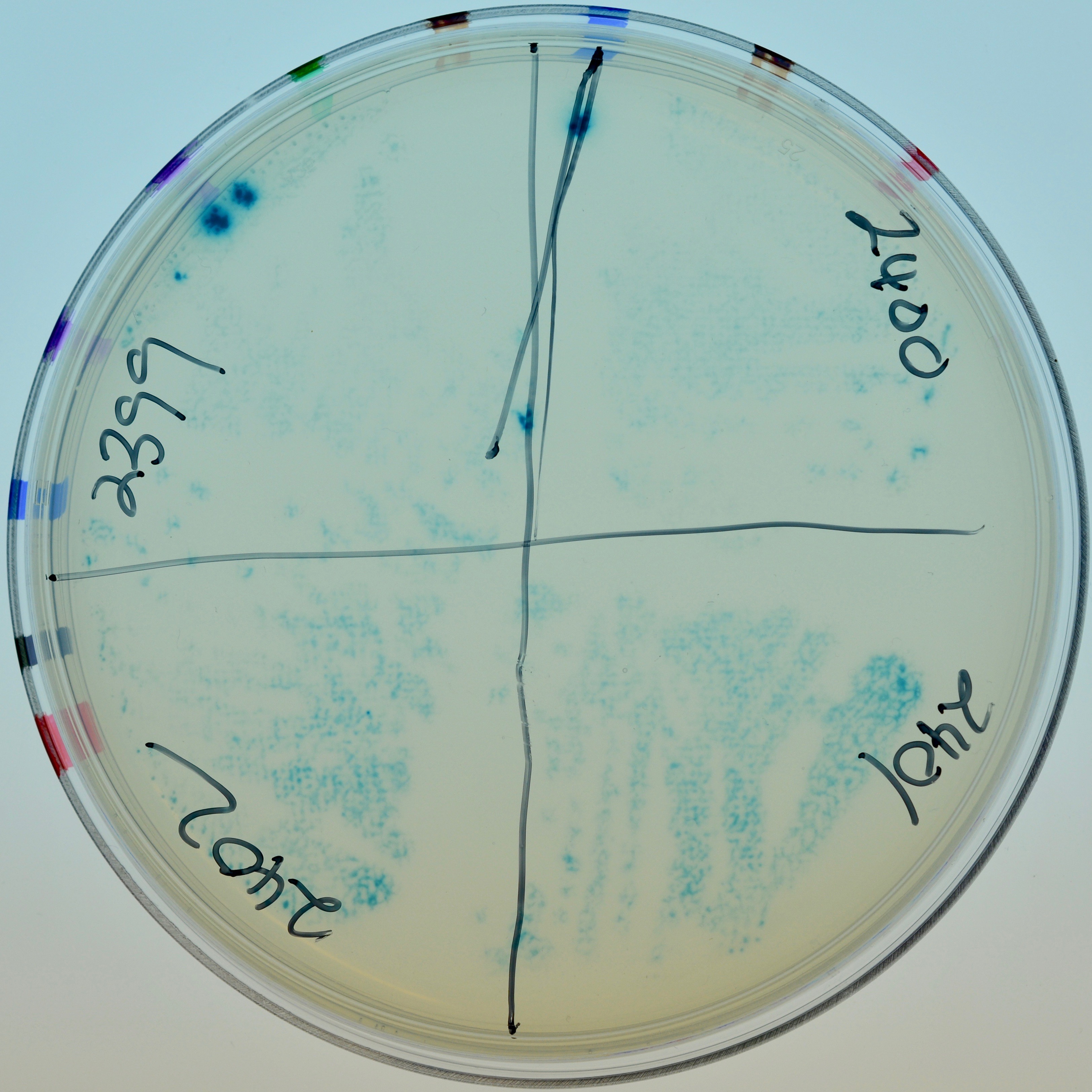

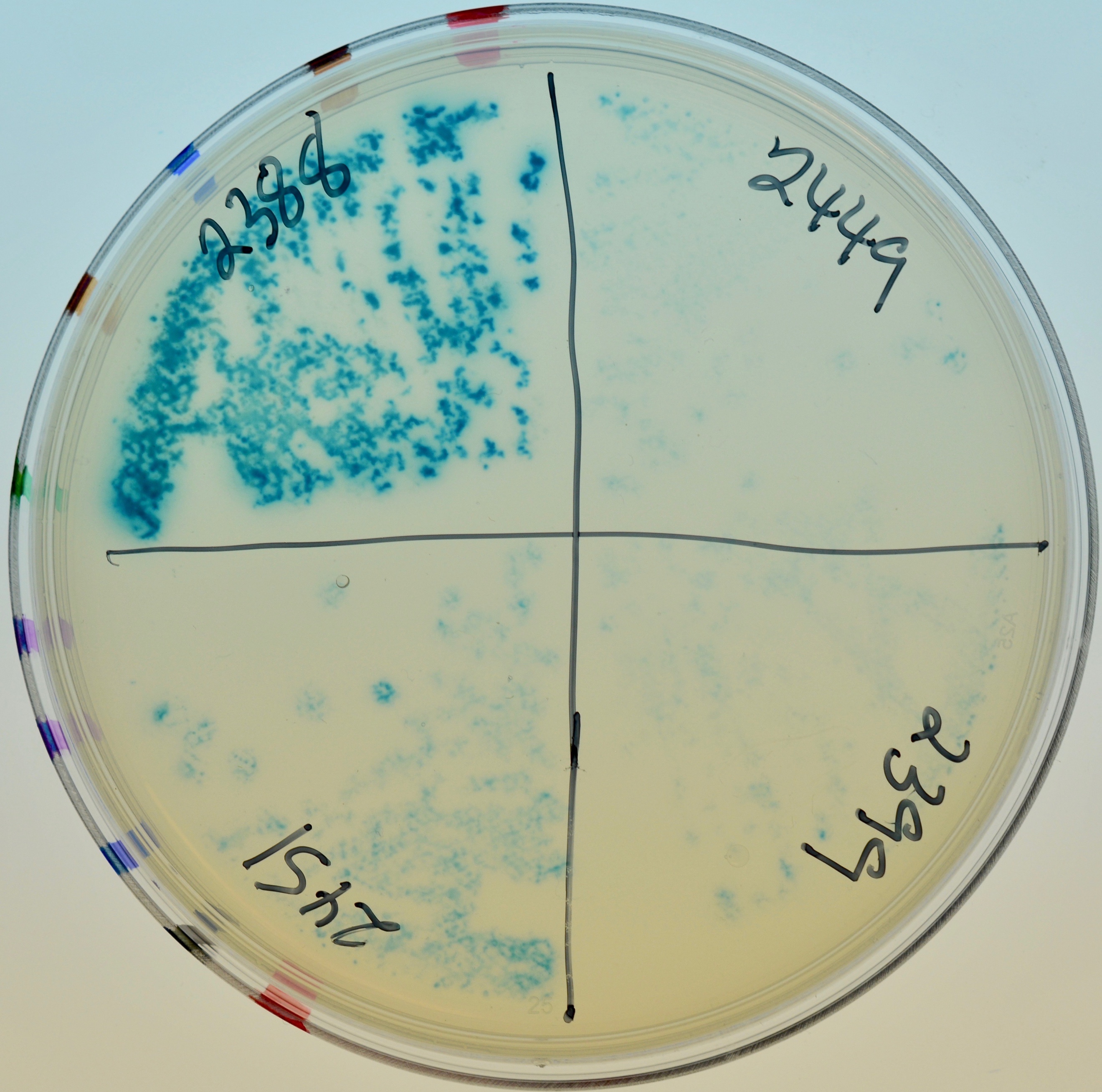

1

Appearance of colonies on minimal maltose with Amp100, Kan30, IPTG and X-gal.

1. From top left, clockwise: LT2387, LT2388, LT2389, LT2390.

2. From top left, clockwise: LT2399, LT2400, LT2401, LT2402.

3. From top left, clockwise: LT2388, LT2449, LT2399, LT2451.

**C. RexB by CI**

| **Strain** | **Relevant Genotype^†^** | **Configurations of hybrid proteins** | **Growth on Minimal Maltose**  **X-gal** |
| --- | --- | --- | --- |
| LT2331 | BTH101[pKT25-*rexB*] [pUT18C-*c*I] | *cya25-rexB*  *cya18-c*I | Yes |
| LT2332 | BTH101[pKNT25-*rexB*] [pUT18C-*c*I] | *rexB-cya25*  *cya18-c*I | Yes |
| LT2383 | BTH101[pKT25-*c*I] [pUT18-*rexB*] | *cya25-c*I  *rexB-cya18* | Yes |
| LT2384 | BTH101[pKNT25-*c*I] [pUT18-*rexB*] | *c*I-*cya25*  *rexB-cya18* | No |
| LT2385 | BTH101[pKT25-*c*I] [pUT18C-*rexB*] | *cya25-c*I  *cya18-rexB* | No |
| LT2386 | BTH101[pKNT25-*c*I] [pUT18C-*rexB*] | *c*I*-cya25*  *cya18-rexB* | No |
| LT2397 | BTH101[pKT25-*rexB*] [pUT18-*c*I] | *cya25-rexB*  *c*I*-cya18* | No |
| LT2398 | BTH101[pKNT25-*rexB*] [pUT18-*c*I] | *rexB-cya25*  *c*I*-cya18* | No |
| LT2446 | LT2331 *nadA::Tn*10 (λ*c*I<>*kan Δ{N-int})* | *cya25-rexB*  *cya18-c*I | Yes |

1

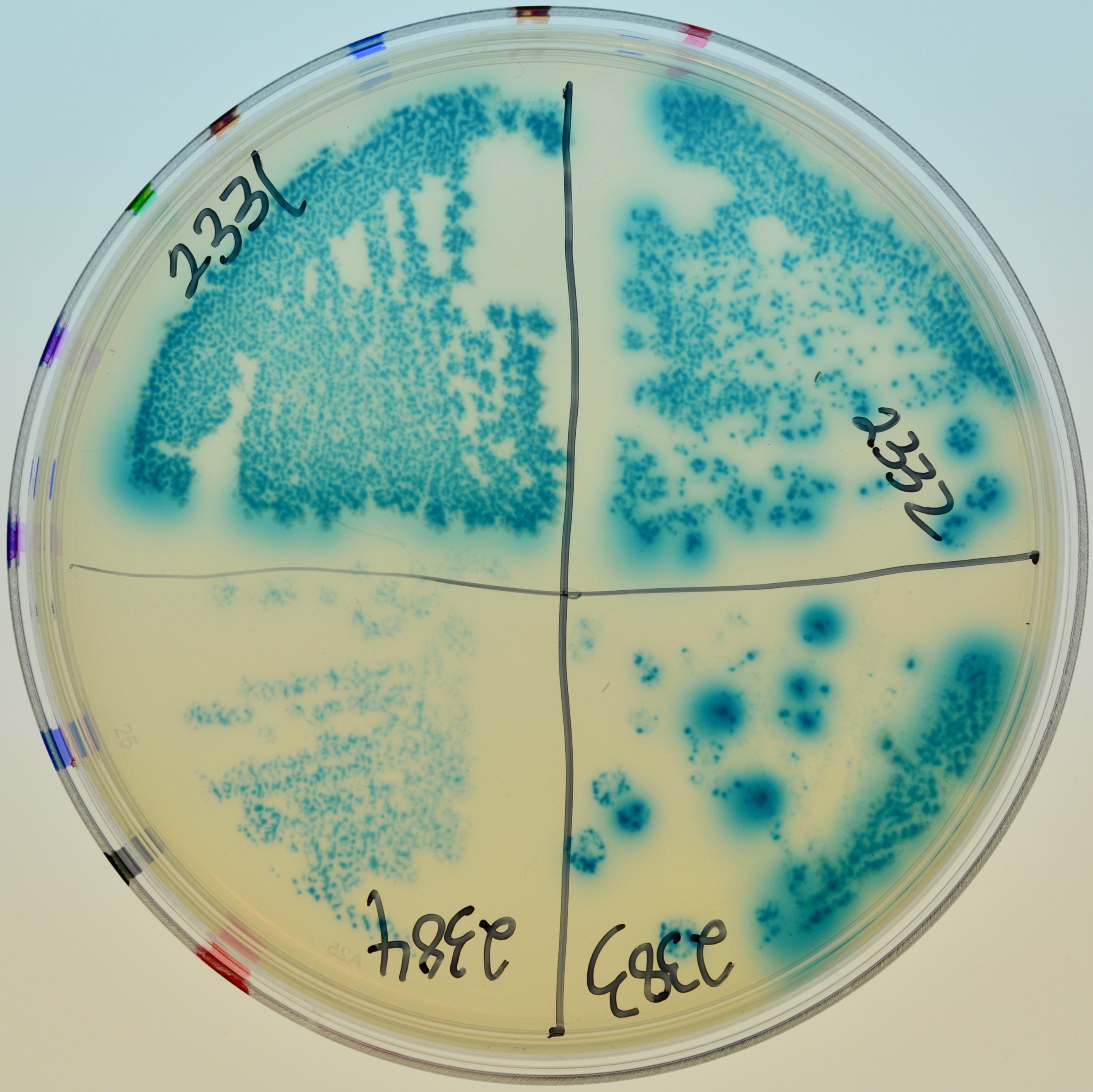

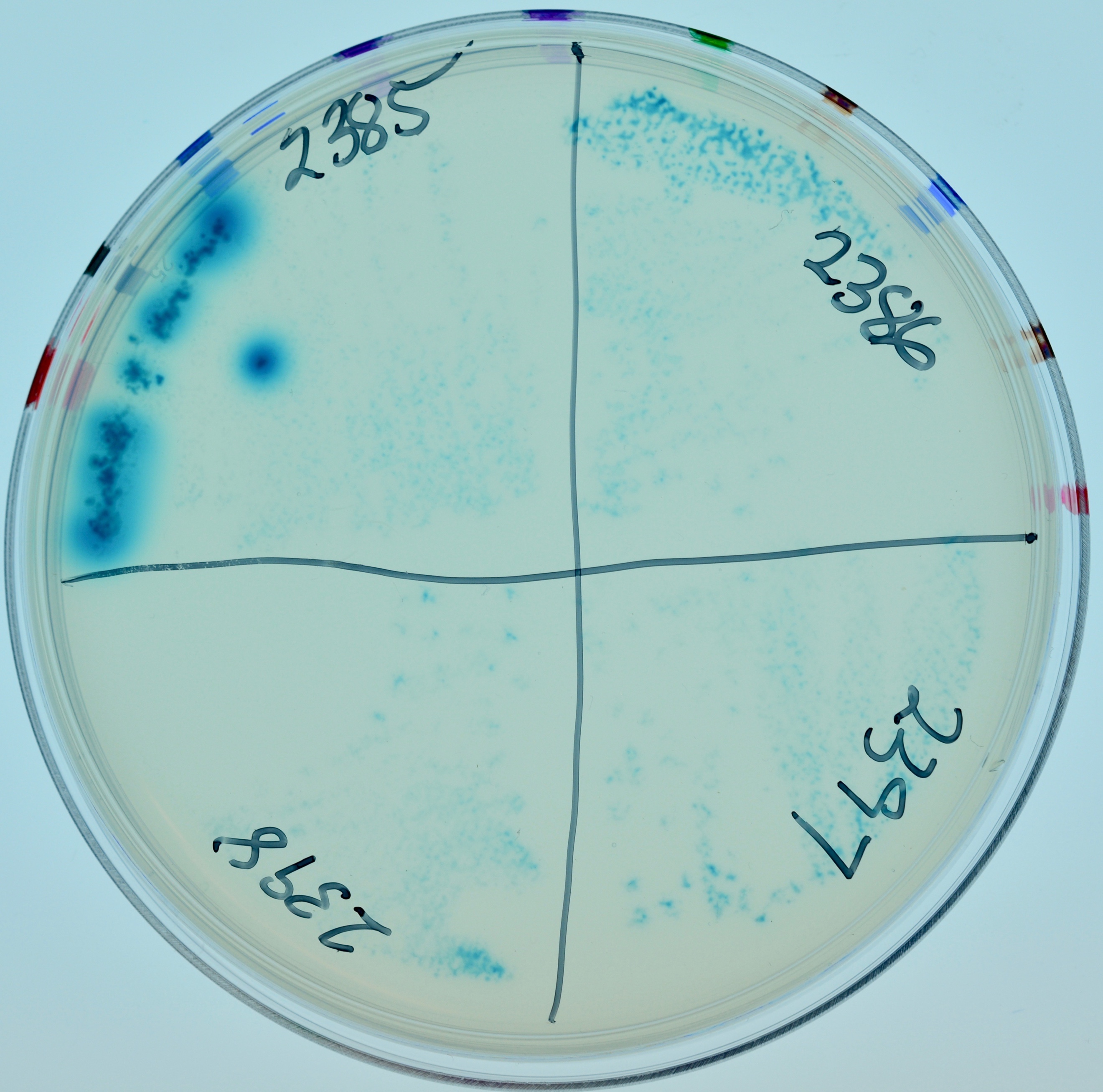

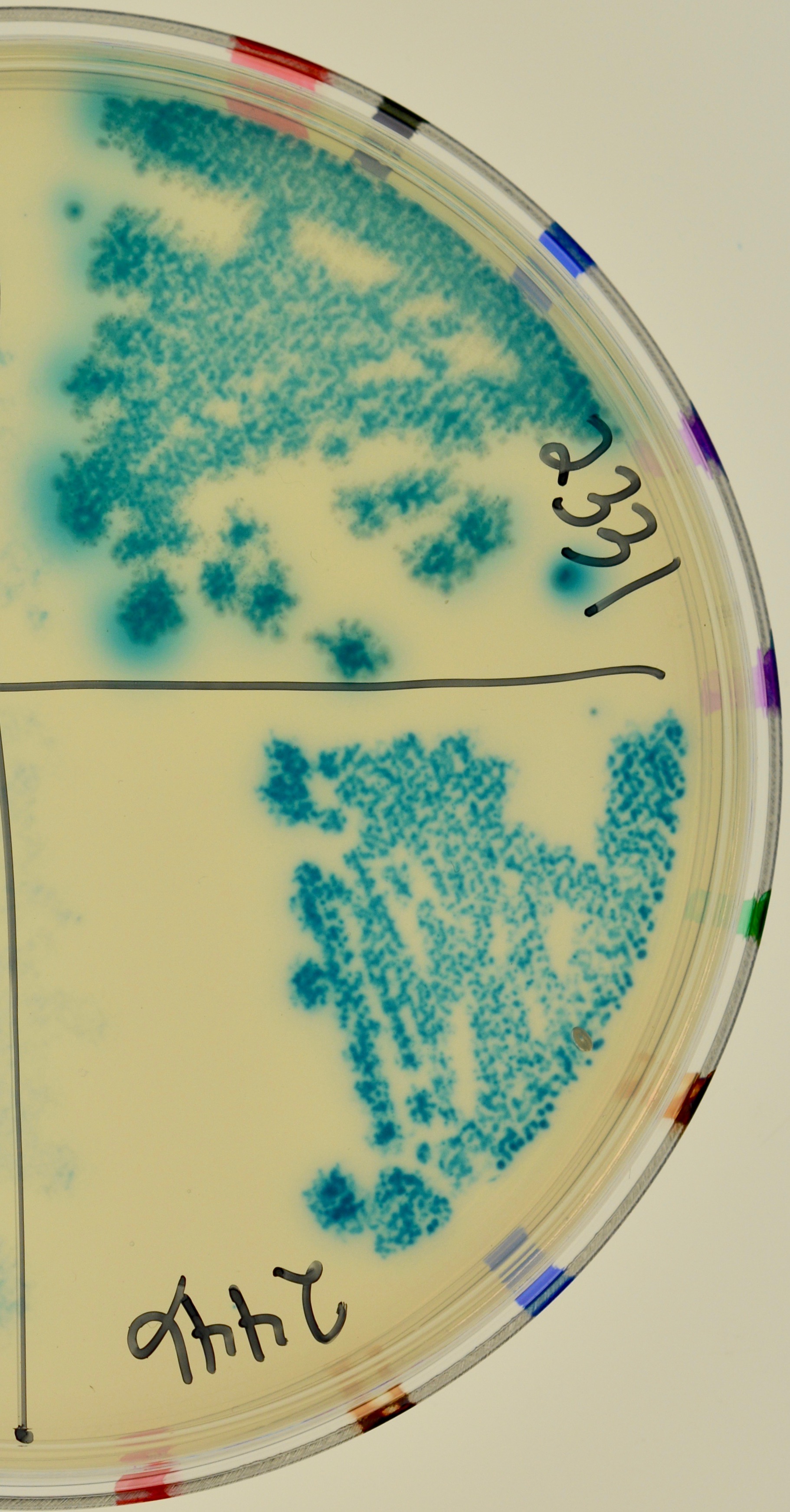

3

2

Appearance of colonies on minimal maltose with Amp100, Kan30, IPTG and X-gal.

1. From top left, clockwise: LT2331, LT2332, LT2383, LT2384.

2. From top left, clockwise: LT2385, LT2386, LT2397, LT2398.

3. Top, LT2331; bottom, LT2446.

**D. RexB by Cro**

| **Strain** | **Relevant Genotype^†^** | **Configurations of hybrid proteins** | **Growth on Minimal Maltose**  **X-gal** |
| --- | --- | --- | --- |
| LT2393 | BTH101[pKT25-*cro*] [pUT18-*rexB*] | *cya25-cro*  *rexB-cya18* | No |
| LT2394 | BTH101[pKT25-*cro*] [pUT18C-*rexB*] | *cya25-cro*  *cya18-rexB* | Yes |
| LT2395 | BTH101[pKT25-*rexB*] [pUT18-*cro*] | *cya25-rexB*  *cro-cya18* | No |
| LT2396 | BTH101[pKNT25-*rexB*] [pUT18-*cro*] | *rexB-cya25*  *cro-cya18* | No |
| LT2405 | BTH101[pKNT25-*cro*] [pUT18-*rexB*] | *cro-cya25*  *rexB-cya18* | No |
| LT2406 | BTH101[pKNT25-*cro*] [pUT18C-*rexB*] | *cro-cya25*  *cya18-rexB* | No |
| LT2407 | BTH101[pKT25-*rexB*] [pUT18C-*cro*] | *cya25-rexB*  *cya18-cro* | No |
| LT2408 | BTH101[pKNT25-*rexB*] [pUT18C-*cro*] | *rexB-cya25*  *cya18-cro* | No |
| LT2452 | LT2407 *nadA::Tn*10 (λ*c*I<>*kan Δ{N-int}*) | *cya25-rexB*  *cya18-cro* | No |

3

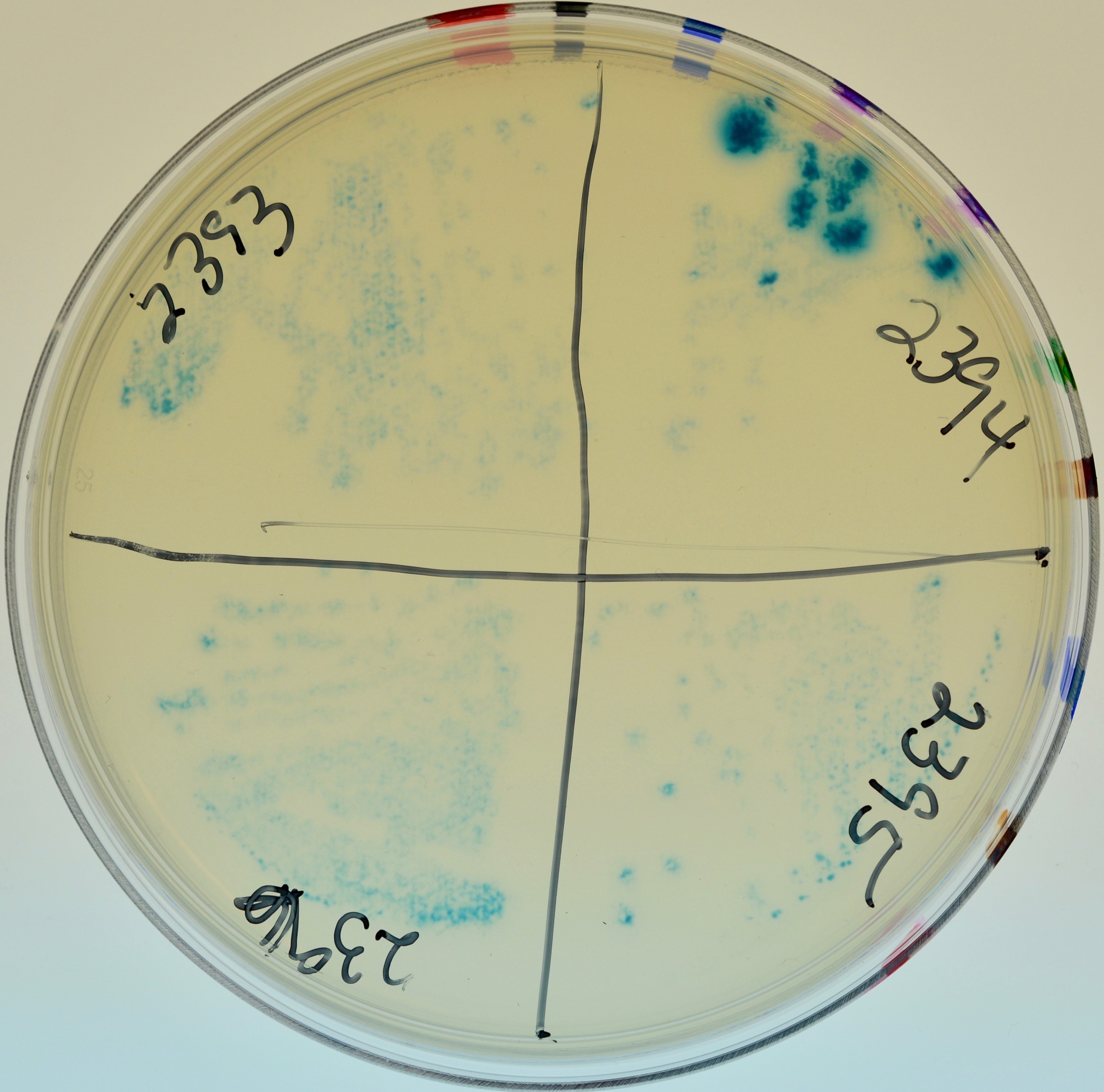

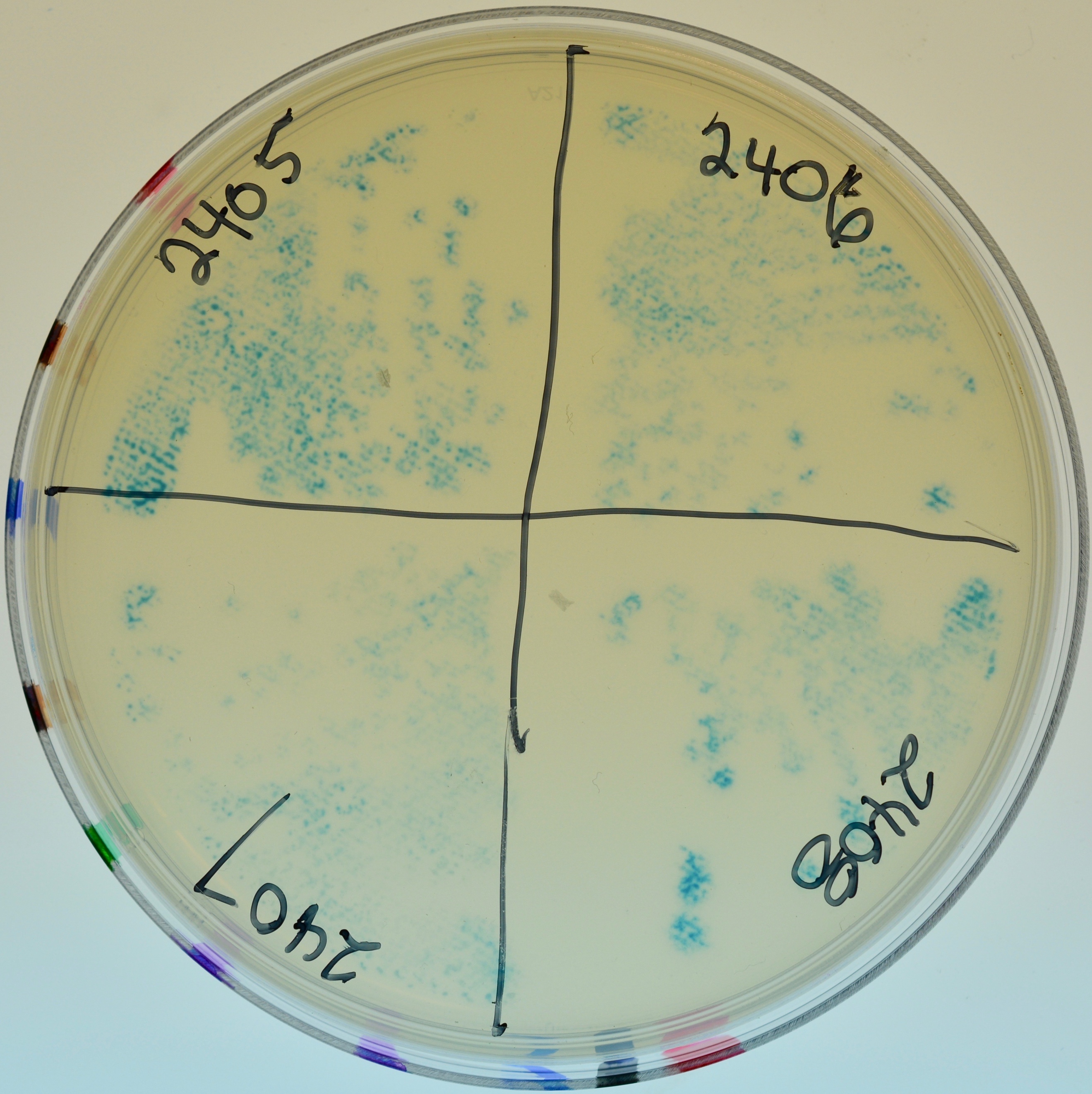

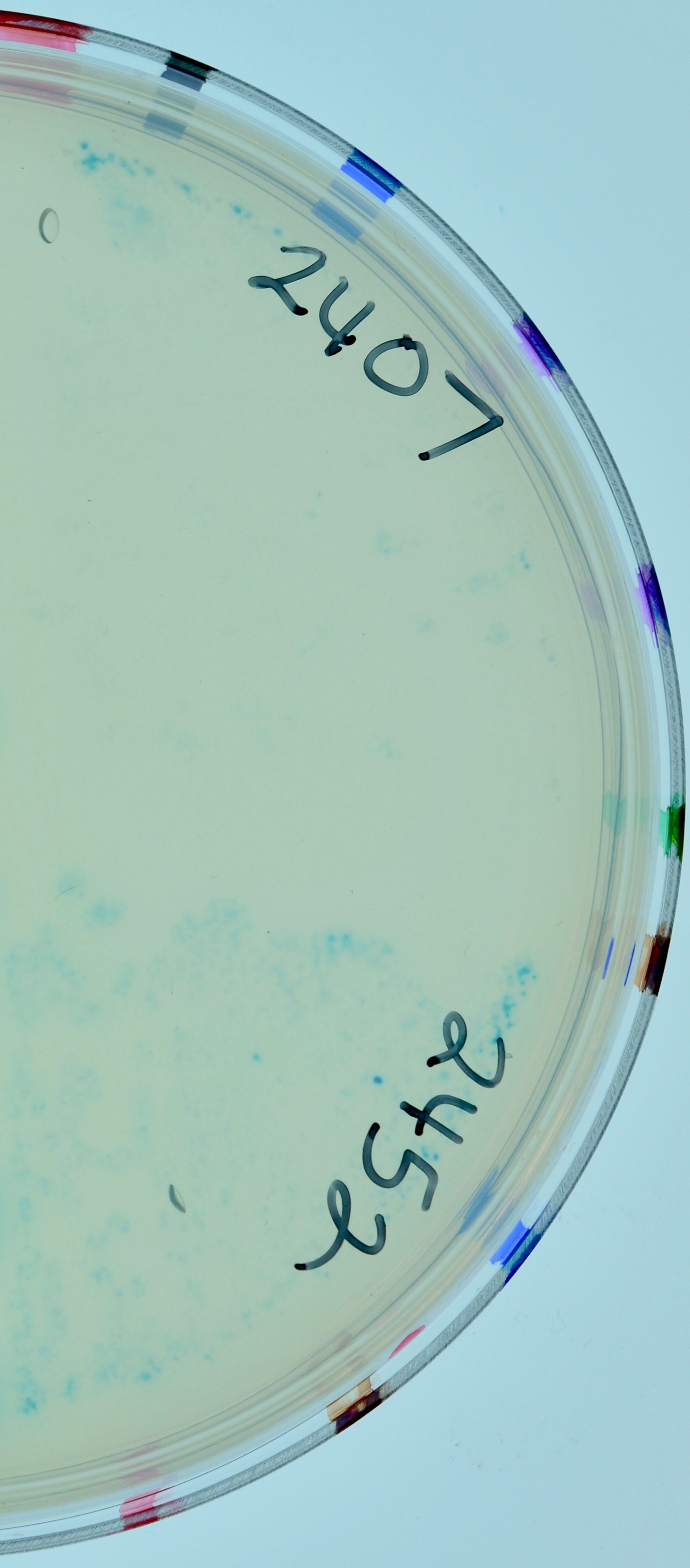

2

1

Appearance of colonies on minimal maltose with Amp100, Kan30, IPTG and X-gal.

1. From top left, clockwise: LT2393, LT2394, LT2395, LT2396.

2. From top left, clockwise: LT2405, LT2406, LT2408, LT2407.

3. Top, LT2407; bottom, LT2452.

**E. cI by Cro**

| **Strain** | **Relevant Genotype^†^** | **Configurations of hybrid proteins** | **Growth on Minimal Maltose**  **X-gal** |
| --- | --- | --- | --- |
| LT2403 | BTH101[pKT25-*c*I] [pUT18-*cro*] | *cya25-c*I  *cro-cya18* | No |
| LT2404 | BTH101[pKNT25-*c*I] [pUT18-*cro*] | *c*I*-cya25*  *cro-cya18* | No |
| LT2409 | BTH101[pKT25-*cro*] [pUT18-*c*I] | *cya25-cro*  *c*I*-cya18* | No |
| LT2410 | BTH101[pKT25-*cro*] [pUT18C-*c*I] | *cya25-cro*  *cya18-c*I | Yes |
| LT2411 | BTH101[pKT25-*c*I] [pUT18C-*cro*] | *cya25-cI*  *cya18-cro* | No |
| LT2412 | BTH101[pKNT25-*c*I] [pUT18C-*cro*] | cI*-cya25*  *cya18-cro* | No |
| LT2413 | BTH101[pKNT25-*cro*] [pUT18-*c*I] | *cro-cya25*  *c*I*-cya18* | No |
| LT2414 | BTH101[pKNT25-*cro*] [pUT18C-*c*I] | *cro-cya25*  *cya18-c*I | No |
| LT2453 | LT2410 *nadA::Tn*10 (λ*c*I<>*kan Δ{N-int}*) | *cya25-cro*  *cya18-c*I | No |
| LT2454 | LT2411 *nadA::Tn*10 (λ*c*I<>*kan Δ{N-int}*) | *cya25-c*I  *cya18-cro* | No |

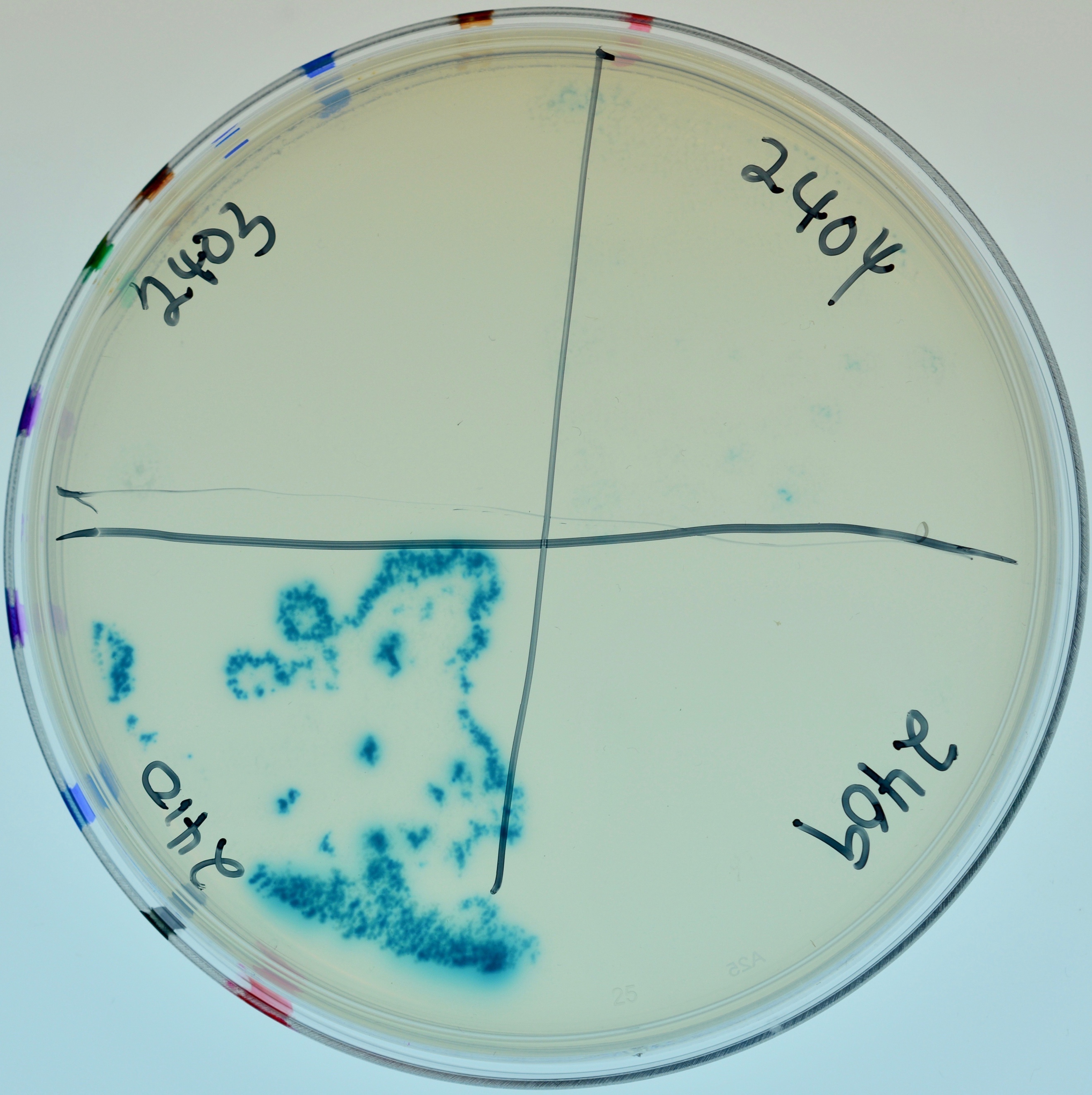

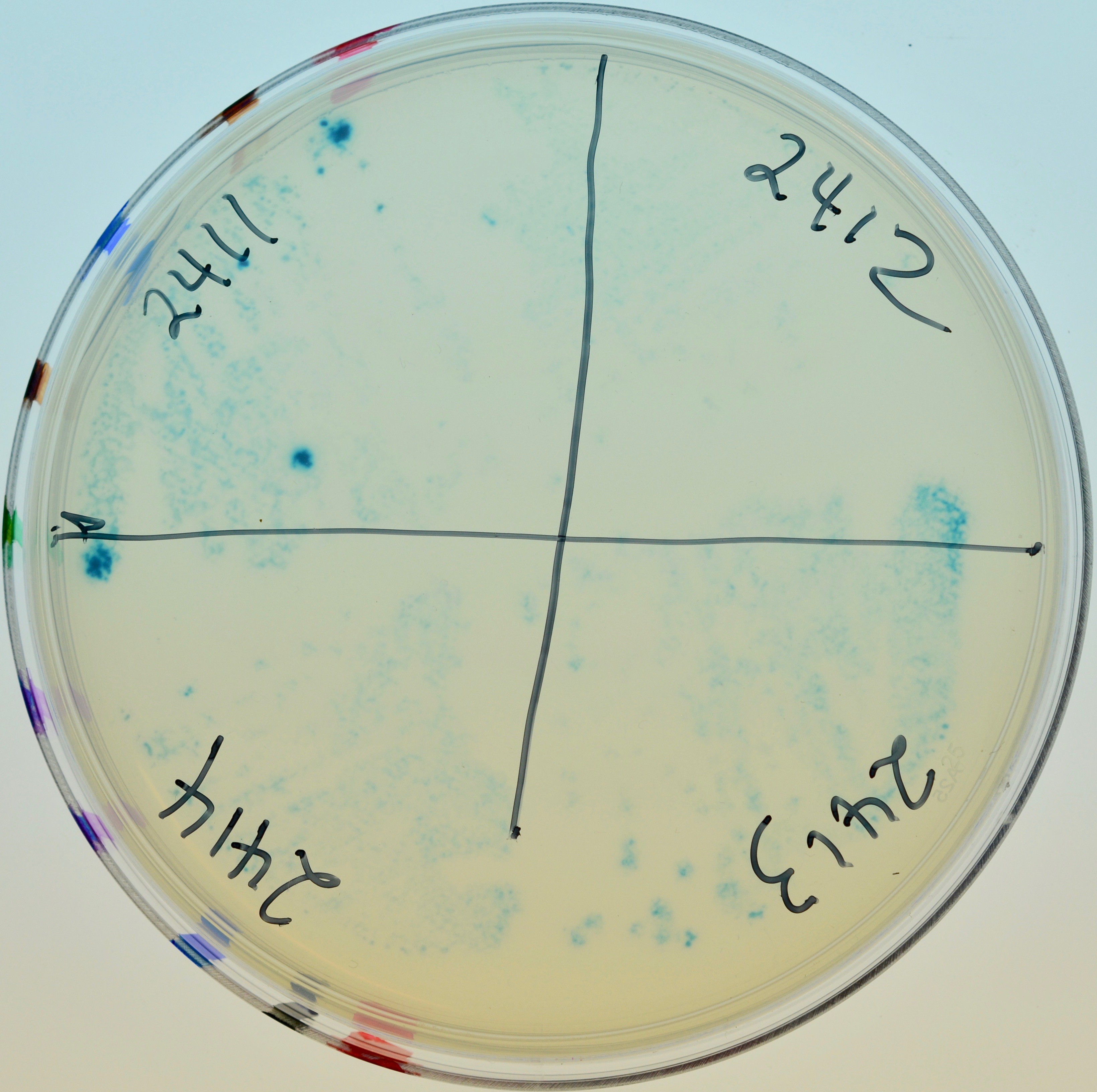

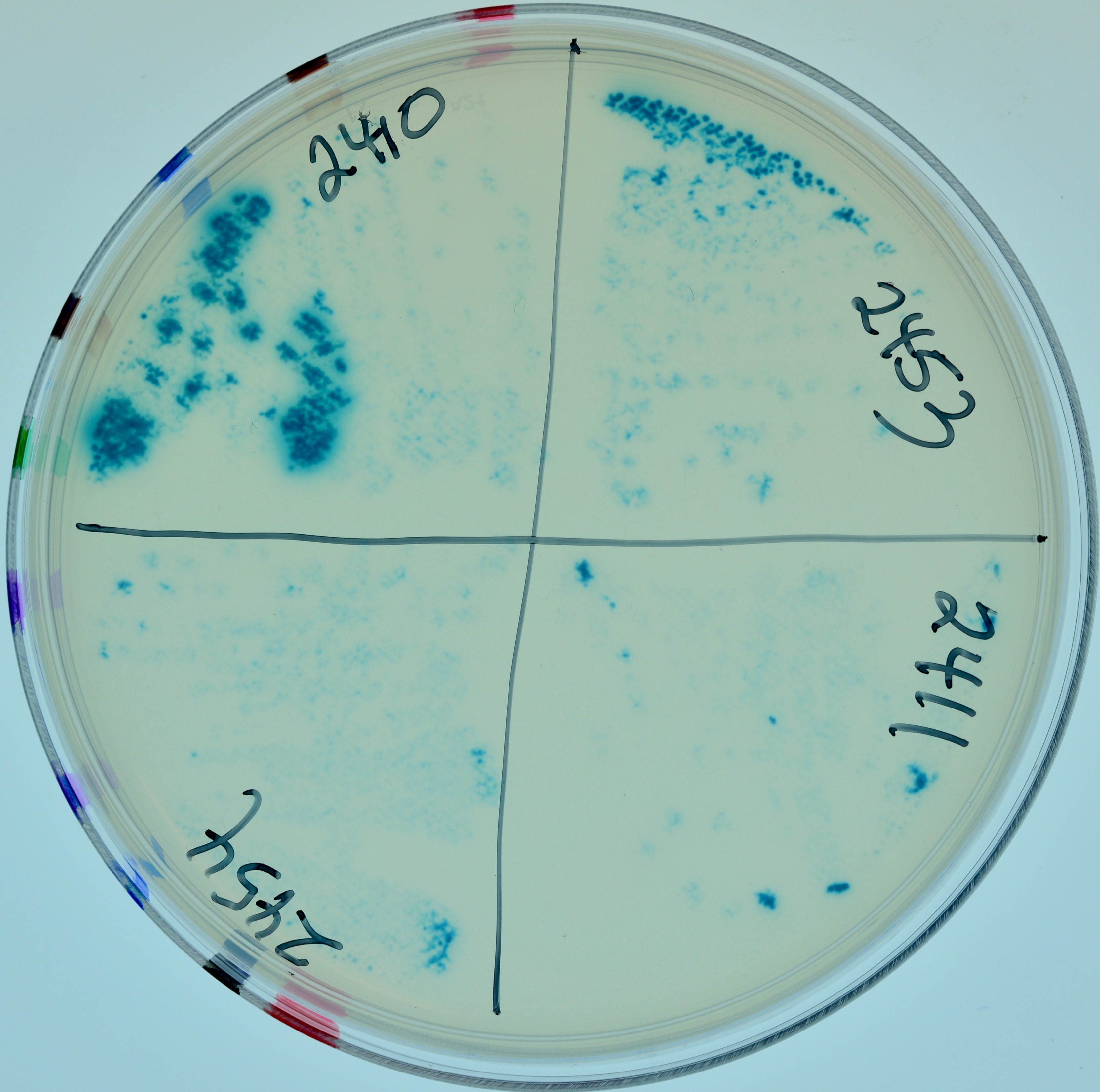

1

2

3

Appearance of colonies on minimal maltose with Amp100, Kan30, IPTG and X-gal.

1. From top left, clockwise: LT2403, LT2404. LT2409, LT2410.

2. From top left, clockwise: LT2411, LT2412, LT2413, LT2414.

3. From top left, clockwise: LT2410, LT2453, LT2411, LT2454.

| **Strain** | **Relevant Genotype^†^** | **Configurations of hybrid proteins** | **Growth on Minimal Maltose**  **X-gal** |
| --- | --- | --- | --- |
| LT2415 | BTH101[pKT25-*c*I] [pUT18C-*c*I] | *cya25-*cI  *cya18-c*I | Yes |
| LT2416 | BTH101[pKT25-*cro*] [pUT18C-*cro*] | *cya25-cro*  *cya18-cro* | Yes |

1

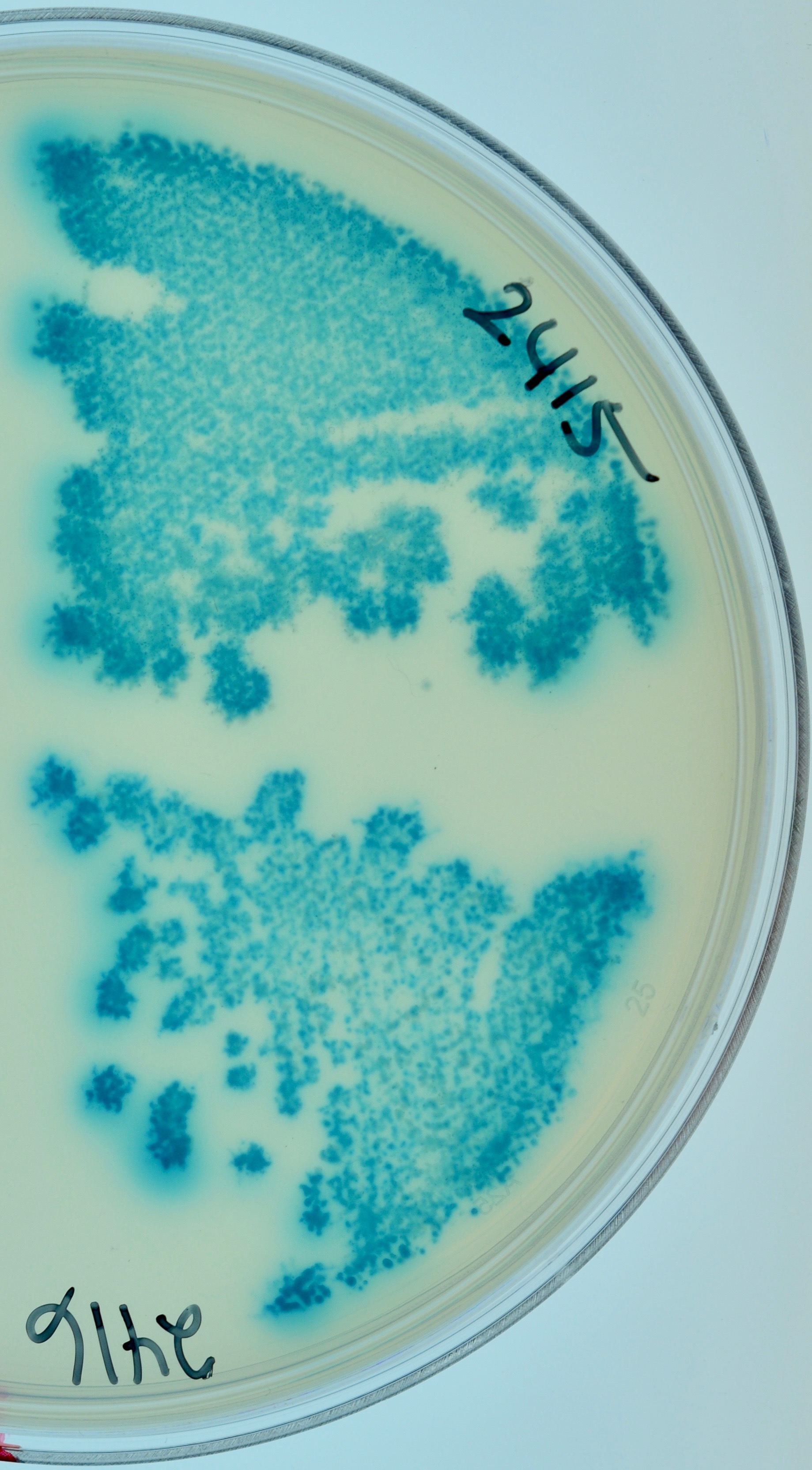

Appearance of colonies on minimal maltose with Amp100, Kan30, IPTG and X-gal.

1. Top, LT2415; bottom, LT2416.

**^†^**Plasmids pKT25 and pKNT25 are low copy and kanamycin resistant (KanR). Plasmids pUT18 and pUT18C are high copy and ampicillin resistant (AmpR).

**Table S3. DNA oligonucleotide sequences^†^.**

| O_R_1-O_R_2 US | 5'-AGATCTA(m^5^C)CGGTAGAGCTT-3' |
| --- | --- |
| O_R_1-O_R_2 LS | 5'-TAAGCTCTA(m^5^C)CGGTAGATC-3' |
| O_R_1-O_R_2 scrambled US | 5'-AGATCTACCGGTAGAGCTT-3' |
| O_R_1-O_R_2 scrambled LS | 5'-TAAGCTCTACCGGTAGATC-3' |
| O_L_1-O_L_2 US | 5'-AGATCTA(m^6^A)CGTTAGAGCTT-3' |
| O_L_1-O_L_2 LS | 5'-TAAGCTCTA(m^6^A)CGTTAGATC-3' |

^†^ ‘us’ and ‘ls’ denote complementary ‘upper strand’ and ‘lower strand’ oligos that were annealed to make double stranded DNA substrates. See Materials and Methods for preparation.

**Table S4. Propensity of the bistable switch to return to the immune state from the nonimmune state (numerical data for graphs in Fig 5A and 5B).**

| **RecA+ strains** | **LT1886**  **RexA^+^ RexB^+^** | **LT1887**  **RexA^-^ RexB^+^** | **LT1891**  **RexA^+^ RexB^-^** | **LT1892**  **RexA^-^RexB^-^** |
| --- | --- | --- | --- | --- |
| % white | 0.9 | 10.0 | 1.6 | 6.2 |
| standard deviation (s.d.) | 0.75 | 6.4 | 0.78 | 4.2 |
| number colonies analyzed | 23 | 23 | 22 | 21 |
| number without white colonies* | 1 | 2 | 2 | 1 |

*discarded from analysis

| ***ΔrecA* strains** | **LT2063**  **RexA^+^ RexB^+^** | **LT2064**  **RexA^-^ RexB^+^** | **LT2065**  **RexA^+^ RexB^-^** | **LT2066**  **RexA^-^RexB^-^** |
| --- | --- | --- | --- | --- |
| % white | 1.7 | 10.1 | 2.7 | 15.0 |
| standard error of the mean (s.e.m.) | 0.72 | 4.6 | 1.78 | 5.5 |
| number colonies analyzed | 17 | 14 | 15 | 14 |
| number without white colonies* | 1 | 1 | 3 | 3 |

*discarded from analysis

**Table S5. Data for Fig. 6: Dependence of phage release on *rexAB* genotype in lysogenic strains growing in liquid culture**

| ***lamB***  **lysogens:** | **LT1684**  **RexA^+^ RexB^+^** | **LT2319**  **RexA^-^ RexB^+^** | **LT2320**  **RexA^+^ RexB^-^** | **LT2321**  **RexA^-^RexB^-^** |
| --- | --- | --- | --- | --- |
| pfu/ml(x10^4^)^†^ | 13.6 | 9.0 | 31.8 | 4.3 |
| standard deviation (s.d.) | 5.0 | 1.1 | 8.3 | 3.1 |

^†^the observed values were multiplied by 10^4^ for simplicity

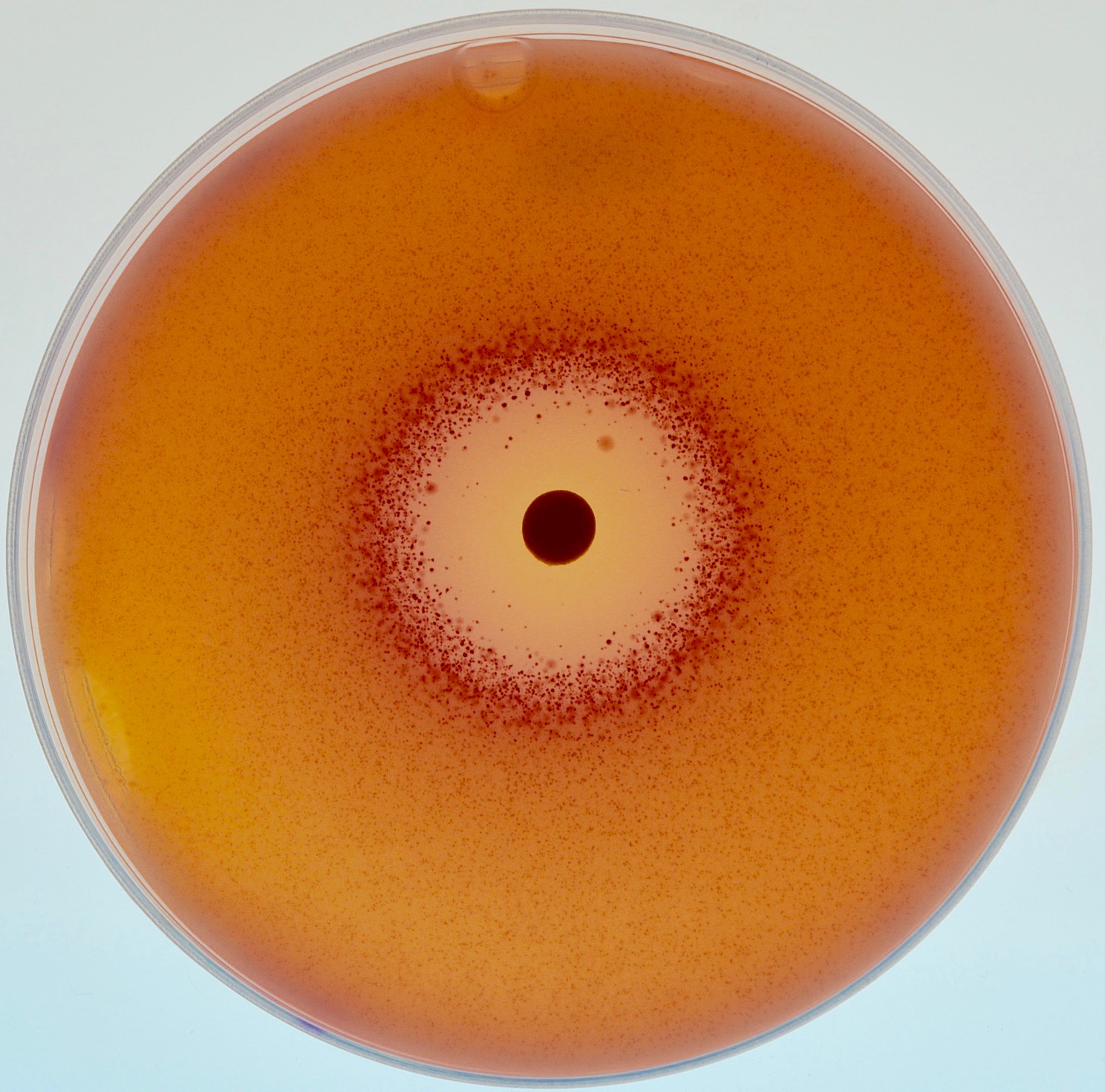

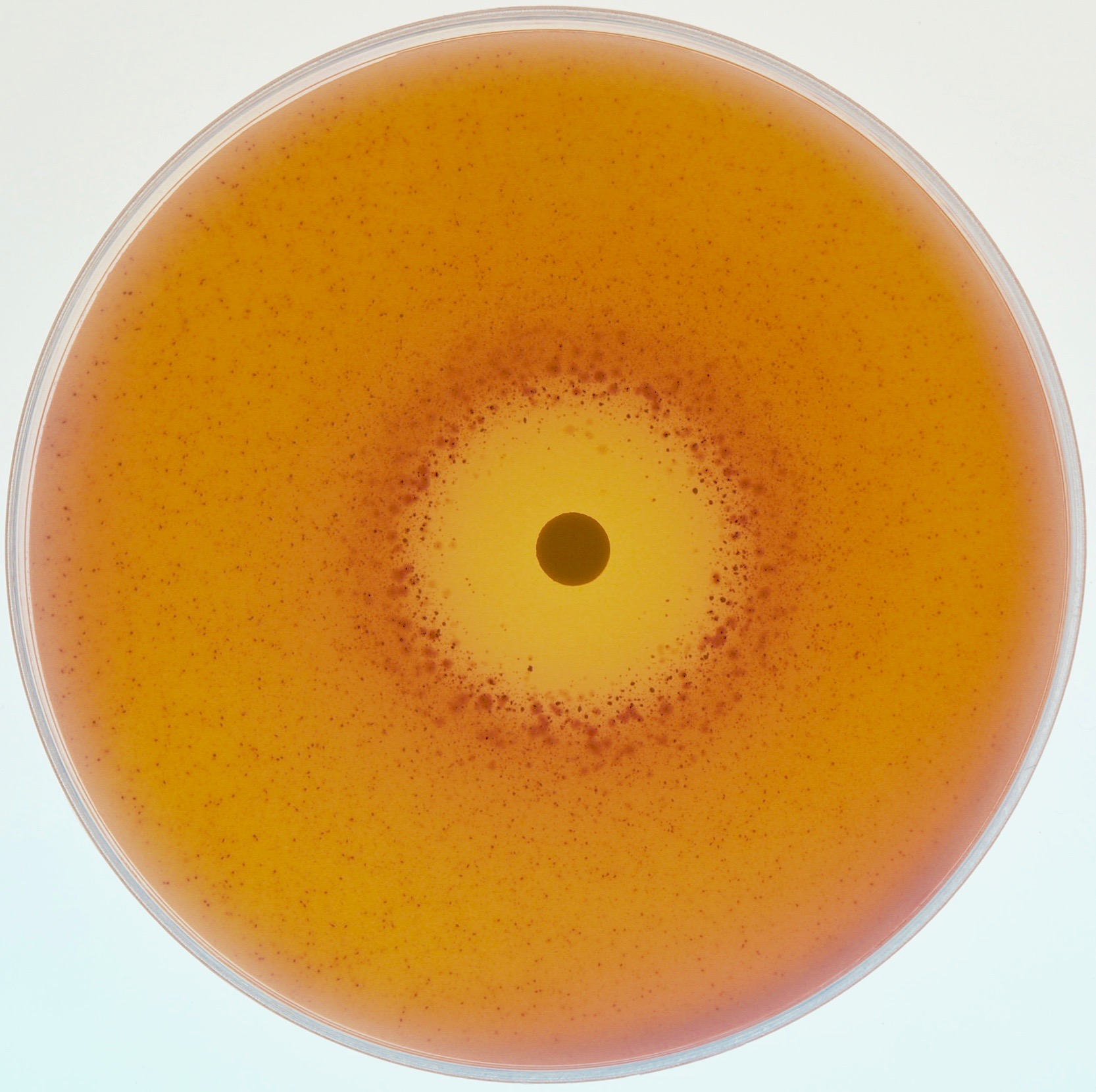

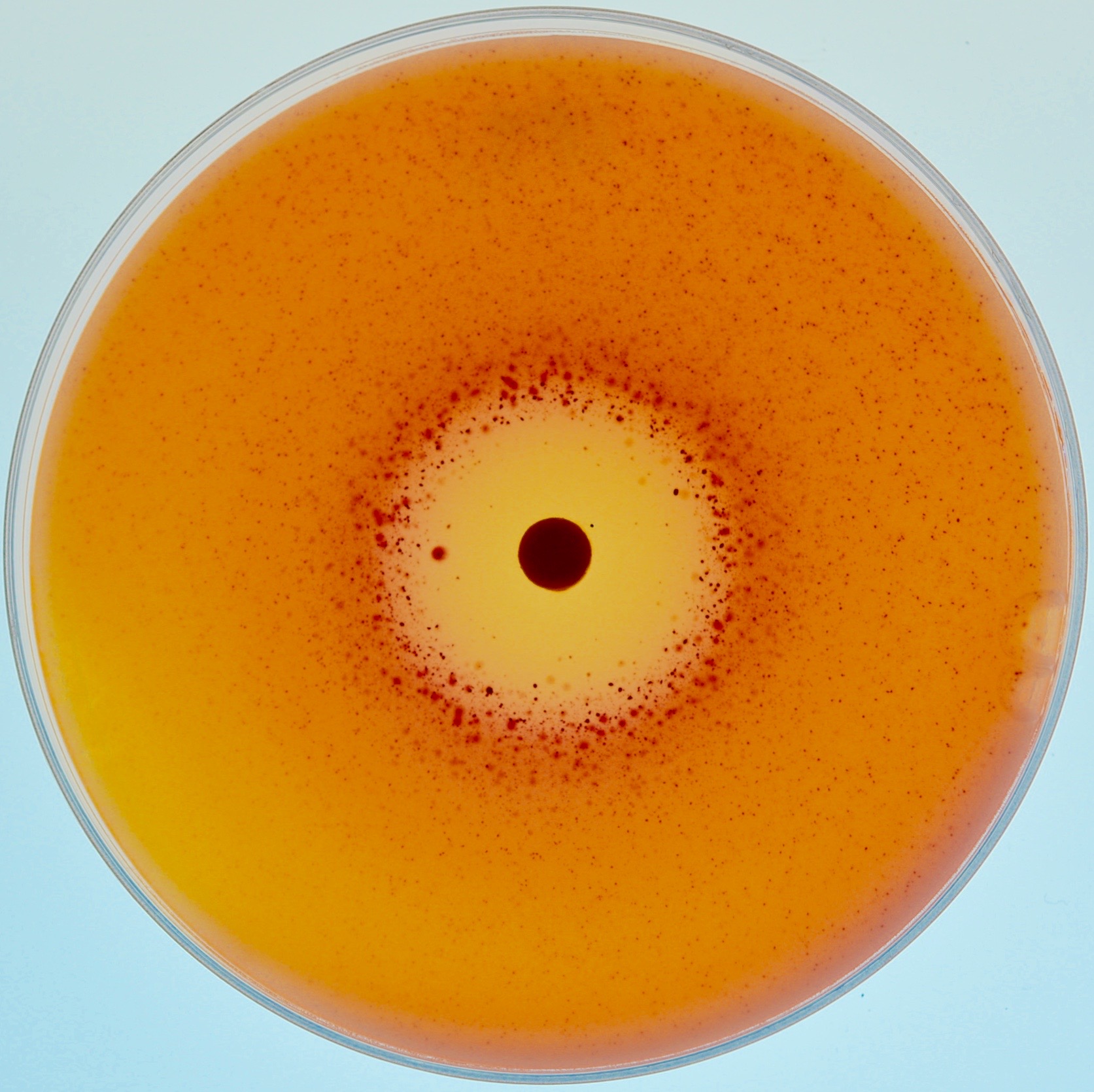

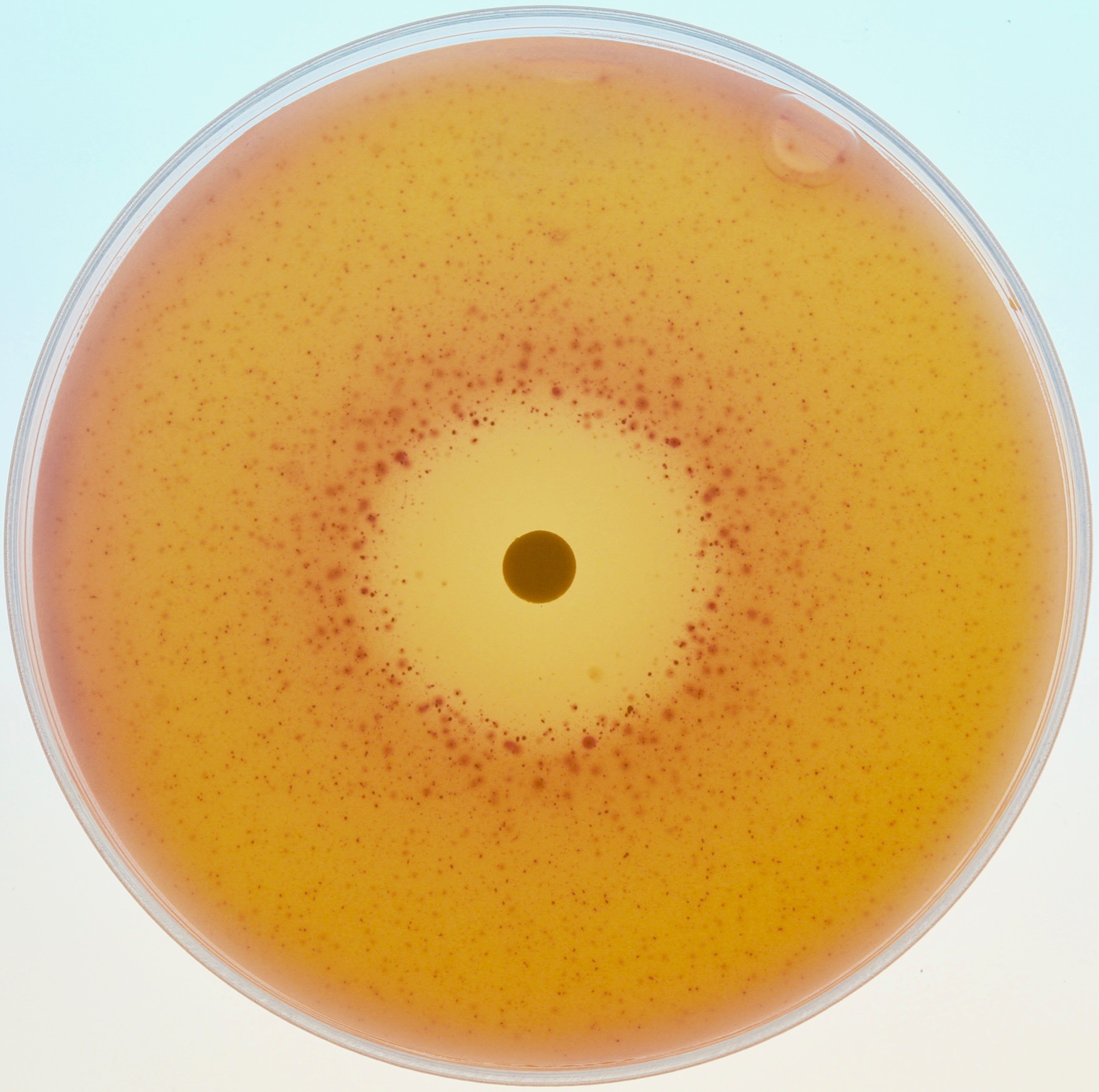

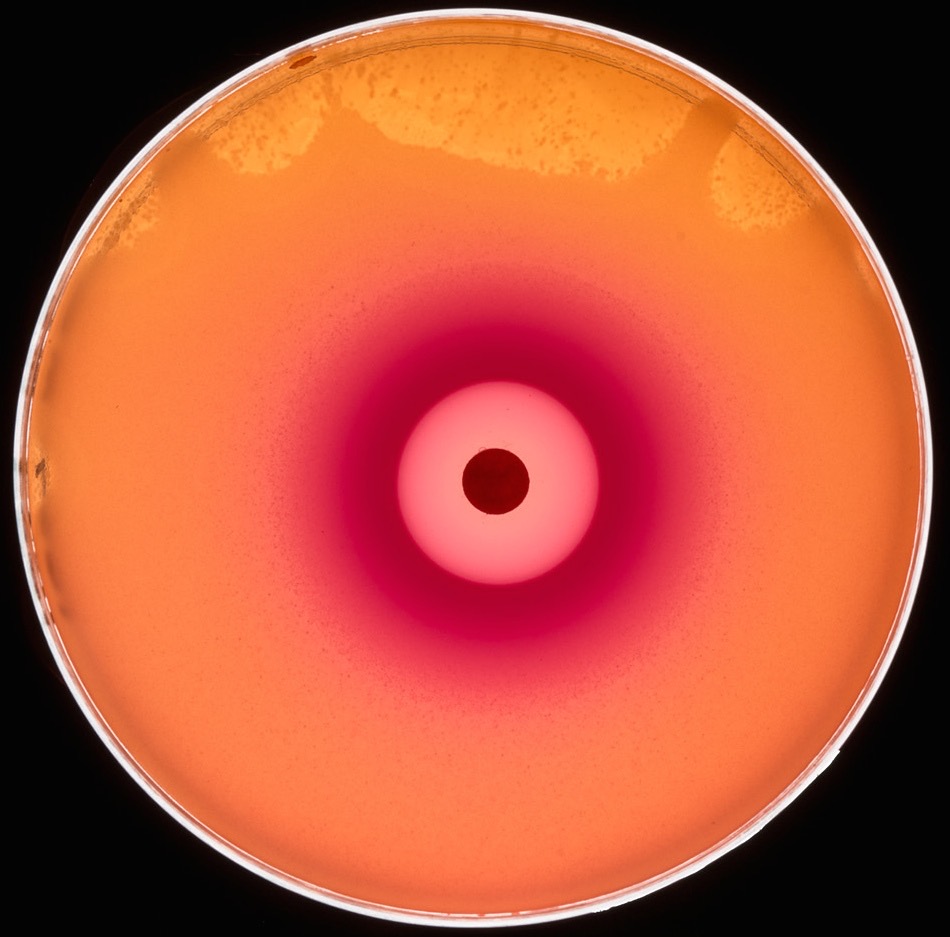

***rexA+ rexB+***

**LT1657**

**LT1886**

***rexA- rexB+***

***rexA- rexB-***

***rexA+ rexB-***

***rexA+ rexB+***

**LT1892**

**LT1891**

**LT1887**

E

A

B

D

C

**Supporting Figure S1. Disk diffusion assays of RecA^+^ Cro^+^ *cI857 ind1* strains challenged with Mitomycin C on MacConkey Lactose indicator media.** Strains **A-D** are used in the experiment shown in Figure 4A; they are RecA^+^ and contain the Cro^+^ genetic construct shown in Figure 3 with a *c*I*857* *ind1* repressor and different *rexA* and *rexB* mutations, as indicated. **E.** LT1657 also has a dual reporter as diagrammed in Figure 3 but is the repressor genotype is *c*I^+^ *ind^+^*. We looked for RecA-dependent switching from the immune to the non-immune state with a disk diffusion assay. A lawn of bacterial cells was poured on the agar, then 8μl of 1μg/ml Mitomycin C was placed on a paper disk at the center of the plates, which were incubated upright at 32ºC. The drug diffuses outward, causing DNA damage and inducing the RecA-dependent SOS response. If the *c*I *ind1* mutation prevents all repressor inactivation, no red ring should be present, since the ring results from RecA*-mediated CI autocleavage and consequent *P_R_-lacZ* expression. However, after two days of incubation at 32ºC, speckled rings appear in response to the drug. Thus, despite being largely uncleavable, the *c*I *ind1* mutation apparently does not entirely prevent RecA*-dependent *c*I repressor cleavage and switching to the non-immune state. Interestingly, the intensity of the rings shows the same dependence on the *rexA* and *rexB* genotypes as is seen in Figure 4. Because of these data, we think that despite the relative tightness of the repressor *ind1* mutation, a low level of RecA-dependent CI repressor inactivation still occurs and is responsible for the differences apparent between Figures 5A and 5B. The *ind1* mutation, E117K, is adjacent to but does not mutate the cleavage site in the CI protein, which is between residues 111 and 112 (Gimble & Sauer, 1985). An alternative explanation, that the red rings result from mutagenesis, since the cells are exposed to a DNA damaging agent seems less likely since mutagenesis would not display RexAB-dependent variation. A similar experiment was attempted with the *recA* mutant strains used in the experiment of Figure 4B, but the bacteria were so sensitive to the drug that extensive cell killing occurred and the plates were not comparable with the RecA^+^ ones. **E.** The Cro^+^ *cI ind^+^* strain LT1657 is provided for comparison; here a strong red ring appears after overnight incubation.

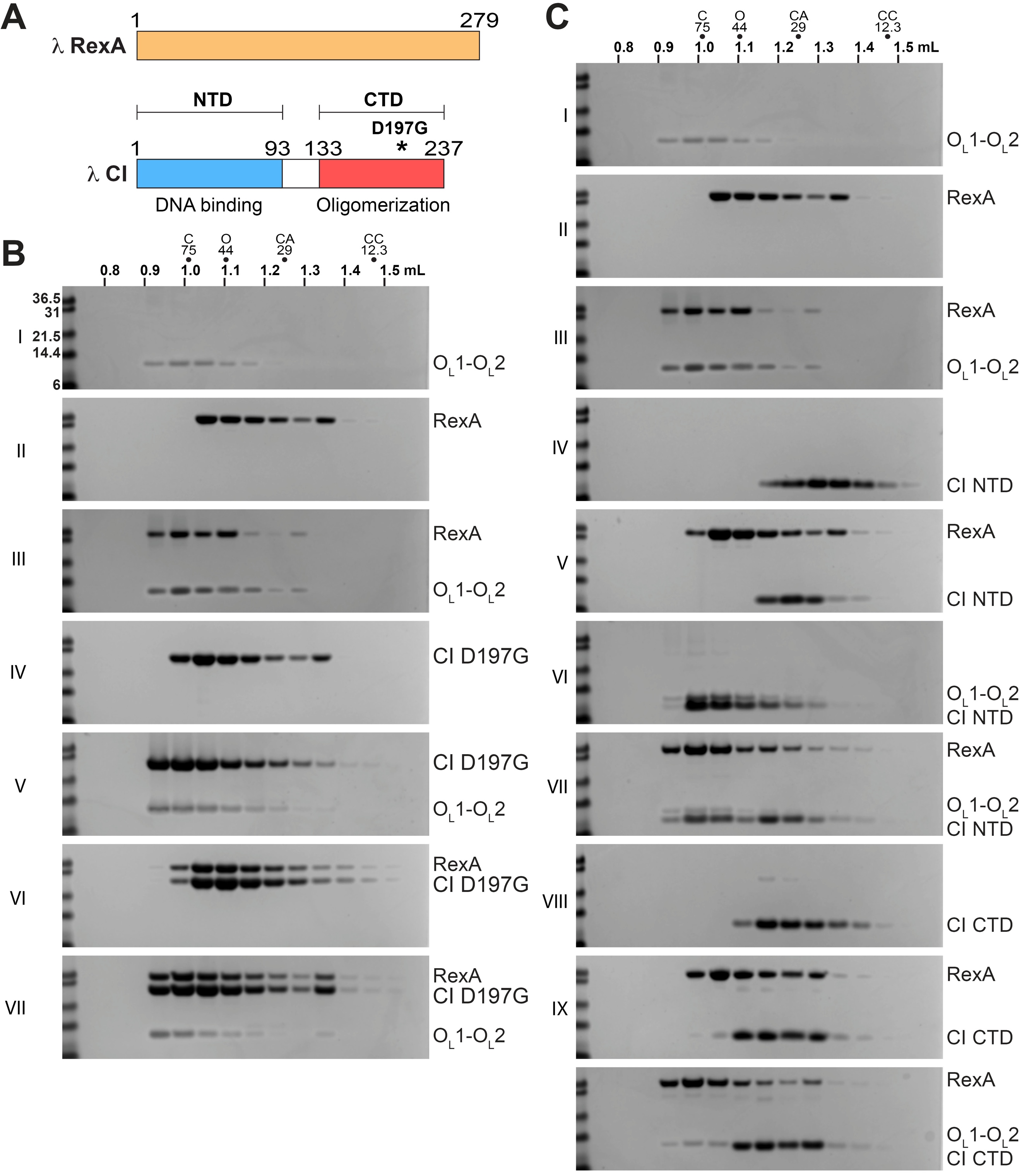
**Supporting** **Figure S2. SEC analysis of CI D197G mutant and truncation constructs.** A. Domain architecture of λ RexA and CI proteins. Boundaries for the CI N-terminal DNA binding domain (NTD, blue) and C-terminal oligomerization domain (CTD, red) truncations are marked along with the relative position of the D197G mutation. B. Analytical SEC analysis of CI D197G protein-protein and protein-DNA interactions. SDS-PAGE gels (silver-stained for DNA and Coomassie-stained for protein) from individual SEC injections are numbered with Roman numerals and shown to visualize shifts in retention volume off of SEC in response to different conditions. Molecular weight standards in kDa are shown in the first lane of each gel with samples labeled on the right. All samples were run on a Superdex 75 PC 3.2 column (GE). A leftward shift of the bands indicates formation of a larger molecular weight species and is associated with complex formation. See Materials and Methods and Table S3 for O_L_1-O_L_2 DNA substrate preparation and oligonucleotide sequences, respectively. C. SEC analysis of protein-protein and protein-DNA interactions of the CI NTD and CTD truncations. Molecular weight standards are the same as in B. Gels I-III are duplicated from B as a reference to allow easier visualization of relative sample shifts.
